# Pangenomic decomposition reveals lineage-specific immune-evasion and virulence architectures in group A *Streptococcus*

**DOI:** 10.64898/2026.05.26.728020

**Authors:** Siddharth M. Chauhan, Jonathan M. Monk, Bernhard O. Palsson, Victor Nizet

## Abstract

*Streptococcus pyogenes* (group A *Streptococcus*, GAS) causes over 700 million infections annually and has resurged globally as a cause of invasive disease. However, the genomic basis by which distinct GAS lineages produce diverse clinical phenotypes remains incompletely understood. Here, we analyzed 399 quality-controlled complete genomes spanning 103 *emm* types and 27 countries and applied non-negative matrix factorization (NMF) to decompose the GAS pangenome into co-inherited gene modules capturing both clonal lineages and mobile elements. This framework revealed that the deepest division in the accessory genome is not defined by *emm* type, but by a 27-gene Sda-1-encoding prophage linked to the hypervirulent M1T1 pandemic clone. This NET-degrading module unites *emm1*, *emm12*, and *emm77* lineages, including fixation in emm77/ST63, extending its distribution beyond classically invasive lineages. Across lineages, virulence determinants segregate into distinct combinations, indicating that invasive disease arises through convergent but mechanistically distinct programs. Consistent with this, highly invasive ST52-*emm28* lacks Sda1 and deploys an alternative repertoire. The decomposition also resolves a gene module corresponding to the emm-pattern D regulon associated with skin tropism, linking accessory genome structure to host niche adaptation. We further identify three sequence-divergent *speC* paralogs on independent prophages. These findings define modular GAS virulence architectures and establish NMF-based pangenome analysis as a framework for genomic surveillance beyond single-locus typing.

**Importance:** Group A *Streptococcus* causes diseases ranging from strep throat to life-threatening flesh-eating infections, with severe cases increasing in recent years and no vaccine currently available. The species is conventionally classified by its M protein gene, but this single-locus system does not capture the extensive genomic diversity driven by prophages that transfer toxin and immune-evasion genes between bacterial lineages. Here, we applied unsupervised machine learning to decompose the pangenome of nearly 400 complete genomes into modules of co-inherited genes. We find that the deepest division among lineages is determined by a prophage carrying a gene that enables evasion of neutrophil extracellular traps, and identify this module at fixation in a lineage not previously known to carry it. We also show that a key toxin gene exists as three independently evolved variants on distinct prophages, and that invasive lineages deploy distinct combinations of virulence factors. This framework enables genomic surveillance based on virulence architecture rather than single-gene typing.

## Introduction

*Streptococcus pyogenes* (commonly called group A *Streptococcus*, GAS) is an exclusively human-adapted pathogen responsible for over 700 million infections and an estimated 500,000 deaths annually ^1^. Its clinical spectrum ranges from superficial pharyngitis and impetigo to life-threatening necrotizing fasciitis, streptococcal toxic shock syndrome, and bacteremia, as well as post-infectious sequelae including acute rheumatic fever and glomerulonephritis ^2^. Despite decades of research, no licensed vaccine exists ^3^, and the molecular basis by which distinct GAS lineages produce diverse disease manifestations remains incompletely understood. Following relaxation of non-pharmaceutical interventions associated with the COVID-19 pandemic, multiple countries reported sharp increases in invasive GAS (iGAS) disease beginning in late 2022, including notable surges in pediatric mortality across Europe ^4,5^, the United Kingdom ^6^, and the United States ^7^. These resurgences, accompanied by expansion of hypervirulent *emm1* sublineages including M1UK ^8^, have renewed urgency for genomic frameworks capable of resolving the mobile virulence architectures that underlie invasive potential across GAS lineages.

GAS population structure is conventionally defined by *emm* typing, which classifies isolates according to sequence variation in the 5’ hypervariable region of the M protein gene ^9,10^. More than 220 *emm* types have been identified ^11^, organized into 48 *emm* clusters reflecting shared binding properties ^12^ and broader *emm* patterns (A-C, D, and E) associated with pharyngeal, skin, and generalist tissue tropisms ^13–15^, respectively. While this framework has substantial epidemiological utility, it captures variation at a single locus and does not account for the extensive accessory genome diversity generated through horizontal gene transfer. In GAS, prophages are the dominant vehicles of genome evolution ^16,17^, and a typical genome harbors 3 to 6 integrated prophages encoding many of the species’ principal virulence determinants, including superantigens (SpeA, SpeC, SpeK), DNases involved in neutrophil extracellular trap (NET) degradation (Sda1, Spd1/Mf2), and phospholipases such as Sla ^2,18^. These prophage complements vary substantially both within and between *emm* types, while Integrative conjugative elements carrying antimicrobial resistance determinants add further complexity ^19^. Together, these observations raise the possibility that clinically important virulence programs may transcend classical *emm*-defined population structure.

Prophage cargo genes are central to GAS host-pathogen interactions and invasive disease biology. In particular, prophage-encoded superantigens and DNases have been implicated in epidemic lineage emergence, immune evasion, and tissue invasion ^20–23^. The hypervirulent M1T1 lineage, including the recently expanded M1UK sublineage ^8^, is associated with prophage-encoded virulence factors including SpeA and the DNase Sda1, which promotes degradation of neutrophil extracellular traps (NETs) during infection. Other prophage-associated virulence factors, including SpeC, SpeK, Spd1/Mf2, and Sla, exhibit variable distributions across lineages and are frequently exchanged through horizontal transfer ^16,17,24^. As a result, GAS disease phenotypes are likely shaped not only by clonal background, but also by distinct combinations of mobile virulence determinants that may cross traditional emm-defined boundaries. However, the broader organizational structure of these co-inherited virulence modules across the GAS pangenome remains poorly resolved.

Large-scale *S. pyogenes* genomic studies have characterized population structure through core genome phylogenetics, emm-based classification, and single-gene analyses of individual virulence determinants ^25–28^. These approaches have provided important insights into lineage emergence and transmission, but they are less suited to resolving the layered organization of the accessory genome, where vertically inherited lineage structure, horizontally transferred mobile elements, and functionally coordinated regulons may coexist within the same chromosomal background. In GAS, this challenge is particularly pronounced because integrated prophages are both major carriers of virulence genes and stable components of long-term lineage evolution. Consequently, clinically important host-interaction programs may be distributed across overlapping combinations of clonal and mobile genetic architectures that are not readily captured by conventional phylogenetic or single-locus frameworks.

Non-negative matrix factorization (NMF) provides a framework for resolving such layered genomic architectures ^29–31^. Applied to a binary gene presence/absence matrix, NMF decomposes the accessory genome into co-occurring gene sets termed “phylons,” which can reflect vertically inherited lineage structure, horizontally transferred mobile elements (mobile phylons, or “mobilons”), or shared functional organization (regulon phylons). This approach has been applied successfully to species including *Escherichia coli* and *Staphylococcus aureus* (^31^, <u>Chauhan et al., in review</u>), where plasmids, genomic islands, and other mobile elements are largely separable from clonal lineage signal. GAS presents a fundamentally different challenge because its accessory genome is dominated by integrated prophages that persist within chromosomal backgrounds over evolutionary timescales while simultaneously encoding many of the species’ principal virulence determinants ^16,20,32,33^.

In GAS, vertical clonal signal and horizontal mobile signal are therefore not merely co-present, but superimposed on the same chromosomal backbone. Integrated prophages typically occupy conserved insertion sites, persist through vertical inheritance across evolutionary timescales, and contribute directly to lineage-specific virulence repertoires ^16,20,32,33^. Consequently, the same category of genes, prophage cargo, simultaneously encodes mobile virulence functions and stable lineage-defining characteristics. Whether matrix decomposition can disentangle these layered inheritance patterns in a highly lysogenized species has not been established. GAS therefore represents a stringent test case for determining whether pangenome decomposition frameworks can resolve the full complexity of bacterial accessory genome organization beyond plasmid- and island-dominated systems.

Here, we apply NMF to a quality-controlled compendium of 399 complete GAS genomes spanning 103 *emm* types, 27 countries, and seven decades of collection. We identify co-inherited genomic modules that resolve both clonal lineage structure and mobile virulence architectures across the GAS pangenome. Through hierarchical exclusive-gene analysis, we show that the primary division among major taxonomic modules is defined not by *emm* type, but by the Sda1-encoding prophage linked to the hypervirulent M1T1 lineage. We further identify Sda1 in *emm77*/ST63, extending the known distribution of this NET-evasion module beyond classically invasive lineages; uncover three sequence-divergent *speC* paralogs carried on independent prophage backbones; and resolve a tissue-tropism regulon associated with skin-adapted lineages. Finally, we show that highly invasive GAS lineages deploy distinct combinations of virulence determinants rather than a shared invasive genotype. Together, these findings establish NMF-based pangenome decomposition as a framework for resolving modular virulence architectures and prophage-driven host-interaction programs in GAS.

## Results

### Assembly of a globally distributed GAS genome compendium

To investigate the accessory genome architecture underlying GAS population structure and virulence, we assembled a curated compendium of complete genomes representing broad geographic, lineage, and tissue site diversity. All publicly available GAS genomes were downloaded from the BV-BRC ^34^ and NCBI RefSeq ^35^ databases (accessed April 2026) and subjected to quality-control procedures adapted from established pangenomic workflows ^31,36,37^. Of 3,267 available assemblies, 399 high-quality, fully circularized (complete) genomes passed filtering criteria and were retained for downstream analysis (**Fig 1a**; see Methods). The curated collection, designated Spyo-GENOMiCUS-400, spans 103 *emm* types from 27 countries and includes isolates recovered from invasive and superficial infection sites including blood, cerebrospinal fluid, bone and joint, skin, and throat/pharyngeal sources (**Fig. 1b, c**).

**Figure 1:**
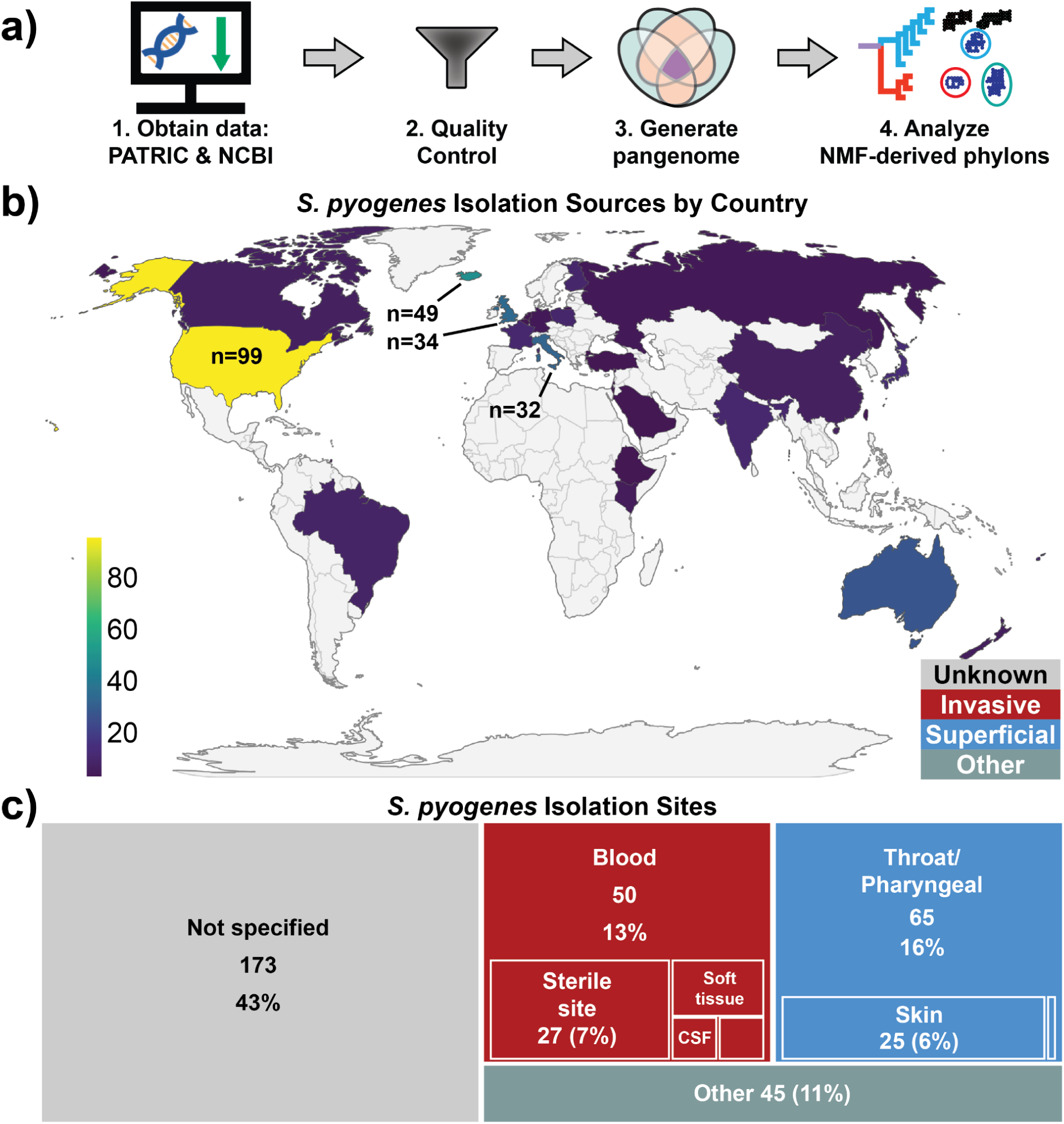
Construction of the Spyo-GENOMiCUS-400 global GAS genome compendium. **(a)** Workflow for assembly of the curated genome collection used for pangenome decomposition and phylon analysis. Complete genomes were obtained from BV-BRC and NCBI RefSeq databases and subjected to quality-control filtering prior to pangenome construction and non-negative matrix factorization (NMF) analysis. ^36^. **(b)** Geographic distribution of the 399 complete genomes included in Spyo-GENOMiCUS-400, representing isolates from 27 countries. The largest contributions originated from the United States (n = 99), Iceland (n = 49), the United Kingdom (n = 34), and Italy (n = 32). **(c)** Distribution of isolation sites across the genome collection, including invasive and superficial infection-associated sources. Blood, throat/pharyngeal, skin, and other sterile-site isolates were broadly represented.

The resulting dataset captures substantial global and clinical diversity within GAS, including lineages associated with pharyngeal colonization, skin infection, and invasive disease. Because prophage acquisition and accessory genome remodeling are major drivers of GAS evolution ^16,17^, we next sought to define how variable gene content partitions across strains and whether co-inherited virulence modules could resolve biologically meaningful lineage structure beyond conventional *emm* typing.

### The GAS pangenome is dominated by a large lineage-variable accessory genome

Following established pangenome partitioning approaches ^36^, we classified genes according to their frequency across the 399 genomes using a cumulative distribution fitted to a double-exponential model, which identifies inflection points separating conserved and lineage-variable gene content (**Fig. 2a,b**, see Methods). The GAS pangenome comprised 8,864 gene families, including 1,292 core genes (present in ≥98.2% of strains), 1,167 accessory genes (present in 8.2-98.2% of strains), and 6,405 rare genes present in <8.2% of strains (**Fig. 2b**). Although rare genes accounted for most gene families in the pangenome, they contributed relatively little to individual genomes, with a median of only 82 rare genes per strain (**Fig. 2c**). In contrast, the accessory genome represented a substantial and broadly distributed component of each genome, with a median of 386 accessory genes per strain, consistent with extensive lineage-specific remodeling through prophage acquisition, recombination, and horizontal gene transfer. In the median genome, core, accessory, and rare genes accounted for approximately 73%, 22%, and 5% of total gene content, respectively. These findings indicate that GAS genomic diversity is structured less by accumulation of highly strain-specific rare genes than by redistribution of a shared accessory gene pool across major lineages.

**Figure 2:**
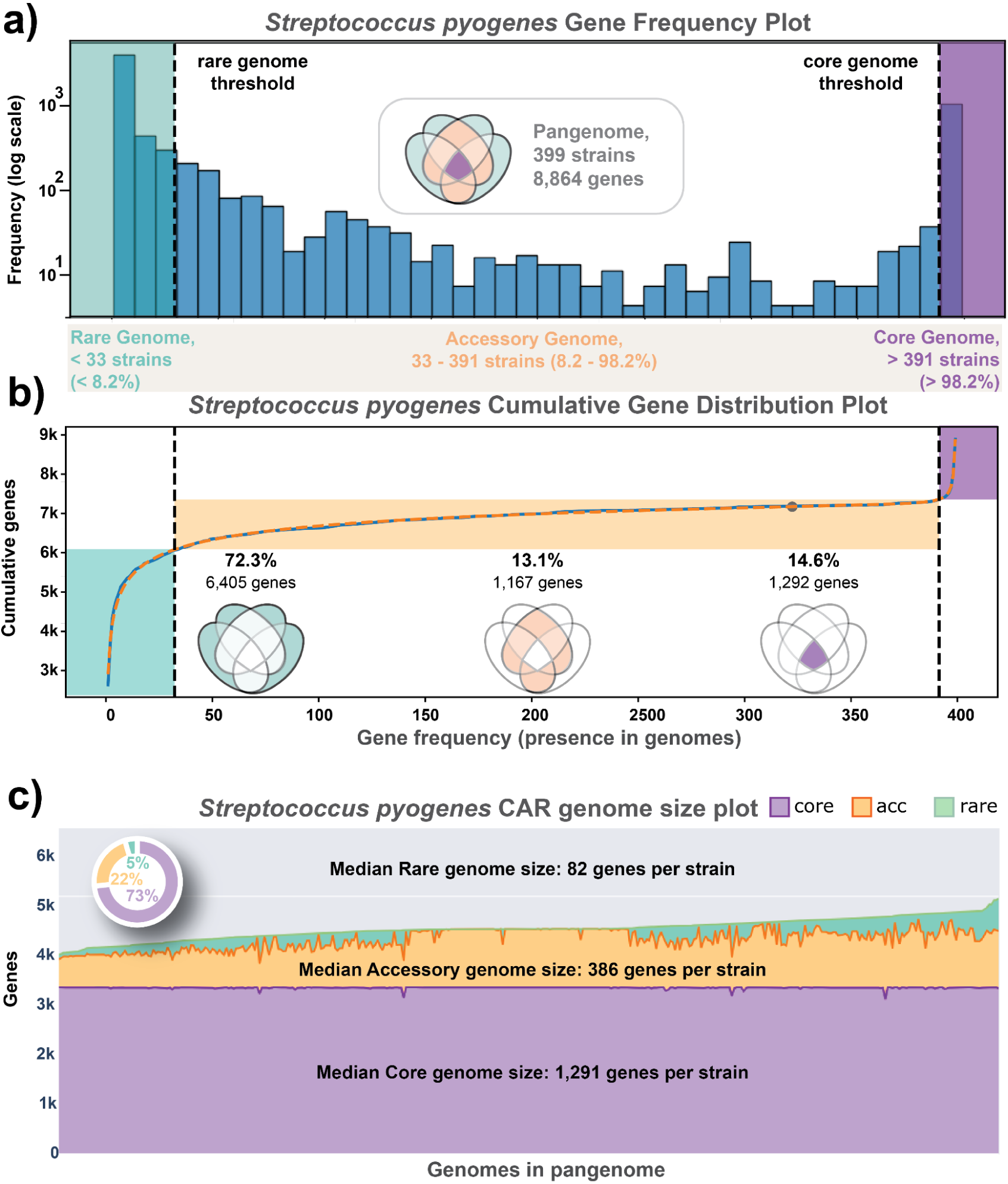
Frequency structure of the GAS pangenome. (**a)** Distribution of gene-family frequencies across the 399 genomes in Spyo-GENOMiCUS-400. (**b)** Cumulative gene-frequency distribution fitted to a double-exponential model ^36^, with median absolute error = 43.41, used to partition the pangenome into core, accessory, and rare gene compartments. The GAS pangenome comprises 8,864 gene families, including 1,292 core genes, 1,167 accessory genes, and 6,405 rare genes. (**c)** Distribution of core, accessory, and rare gene content across individual genomes. The median genome contains approximately 73% core genes, 22% accessory genes, and 5% rare genes.

### Conserved virulence determinants reside within a flexible GAS core genome

Of the 1,292 core genes, 747 (57.8%) were present in all 399 genomes (the omnipresent-core) (Chauhan et al., in review), while the remaining 545 constituted the soft-core ^38^. The omnipresent core encoded the central cellular machinery, major regulators (CovR, CodY, RelA, VicK, LiaF) ^39–44^, and essential cell-division and envelope biosynthesis pathways apparatus ^45^. Among established virulence determinants, only the NADase *nga* ^46,47^ was fully omnipresent. In contrast, many canonical GAS virulence factors occupied the soft core, including the streptolysin S-associated sag operon, speB, ska, scpA, spd, cfa, and dlt loci ^2,48^, each present in 394-397 of 399 genomes. Likewise, the hyaluronic-acid capsule operon (*hasA-C*; 83–88%) ^49–51^, *slo* ^46,52^, and the global virulence regulator *mga* ^13^ (97% each) fell below strict-core thresholds because of lineage-restricted absence or sequence divergence. Thus, even classical GAS virulence programs exhibit substantial evolutionary plasticity, with conserved pathogenic functions maintained through partially overlapping lineage-specific architectures rather than strict universal conservation. Additional lineage-restricted structural variants identified within the core-accessory boundary are described in **Supplementary Note S1** (**Tables S1–S2, Figs. S1–S3**).

### Accessory genome decomposition identifies lineage-specific and mobile virulence modules

To identify the co-inherited gene modules underlying GAS lineage structure, we applied non-negative matrix factorization (NMF) to the binary presence/absence matrix of the 1,167 accessory genes across the 399 genomes (**Fig. 3a**). NMF decomposes this matrix into an **L** matrix defining co-occurring gene sets (“phylons”) and an **A** matrix defining genome-specific phylon affinities (see **Methods**) ^29,31^. Because each genome is reconstructed as a non-negative combination of pylons, the framework captures the modular organization by which GAS genomes are assembled through vertical inheritance layered with prophage acquisition and horizontal gene transfer ^25,26,31^ (**Fig. 3b**).

**Figure 3:**
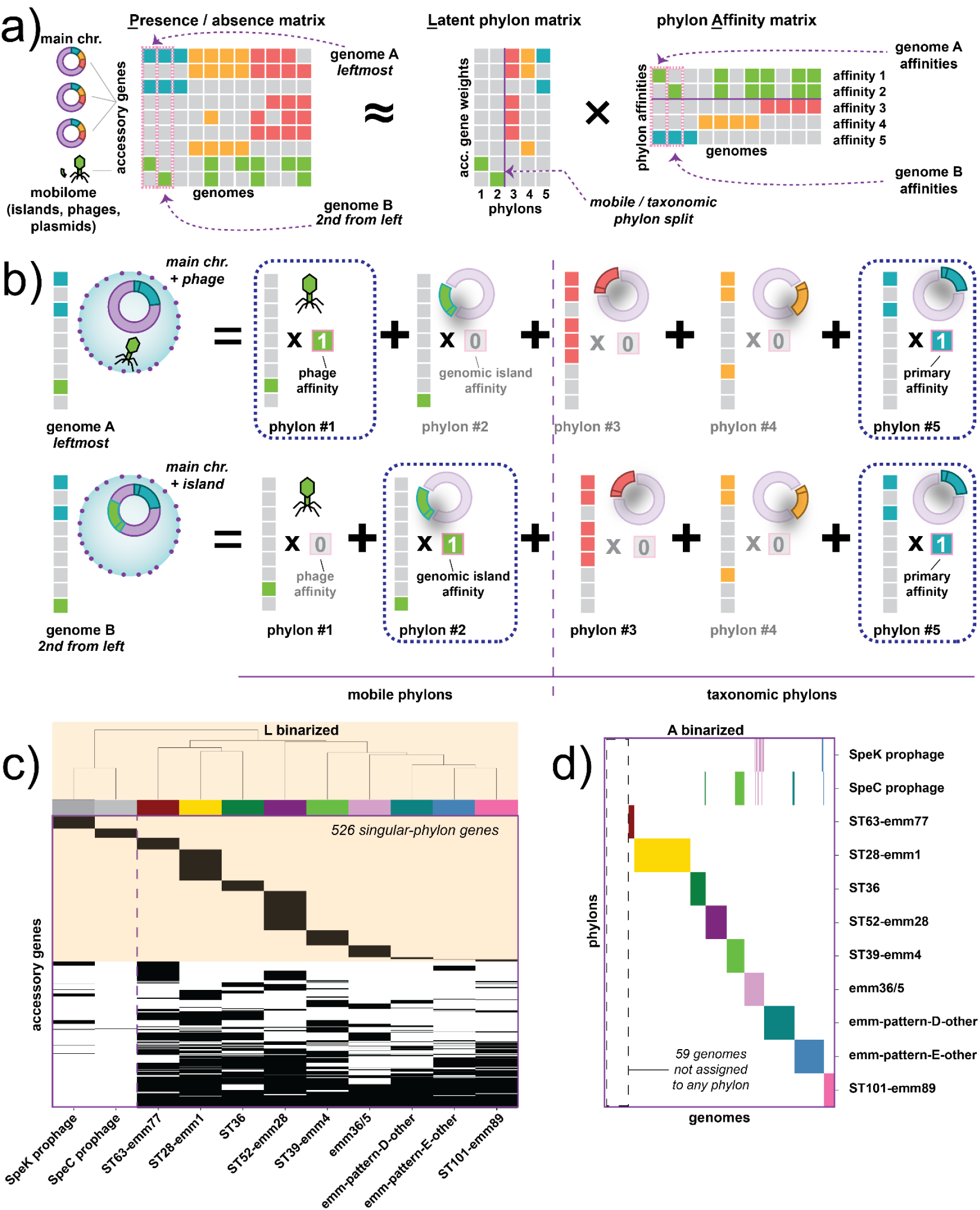
NMF decomposition identifies lineage-specific and mobile accessory genome modules. **(a)** Schematic overview of non-negative matrix factorization (NMF) applied to the accessory gene presence/absence matrix. The matrix is decomposed into an **L** matrix defining co-occurring gene modules (“phylons”) and an **A** matrix defining genome-specific phylon affinities. (**b)** Conceptual model illustrating how GAS genomes are reconstructed as additive combinations of lineage-associated and mobile-element-associated phylons. **(c)** Hierarchical clustering of binarized phylon gene content identifies discrete taxonomic phylons and mobile prophage-associated mobilons. **(d)** Genome-level phylon assignments across the 399-genome dataset showing mutually exclusive taxonomic phylons and sporadic mobilon carriage.

The decomposition identified 11 reproducible pylons that segregated into two biologically distinct classes. (**Fig. 3c,d**). Nine represented taxonomic phylons corresponding to lineage-specific accessory gene repertoires, whereas two represented mobile phylons (“mobilons”) corresponding to independently circulating prophage elements. Most genomes (302/399, 76%) mapped to a single taxonomic phylon, while 38 genomes (10%) carried an additional mobilon. Notably, no genome simultaneously carried two taxonomic phylons, indicating that despite extensive prophage flux, lineage-specific accessory genome structure remains strongly partitioned across the species. Fifty-nine genomes (15%) were not assigned to a phylon, consistent with sparse representation of rare lineages whose accessory gene content fell below the decomposition threshold.

The resulting phylons corresponded closely to known biological groupings within GAS. Seven represented dominant emm-associated lineages (ST28-emm1, ST52-emm28, ST39-emm4, emm3/6/5, ST36, ST63-emm77, and ST101-emm89), two represented composite groups enriched for emm-pattern D or E regulon architectures (14), and two represented independently circulating superantigen-associated prophages. Together, these findings indicate that the GAS accessory genome is organized into a limited number of discrete lineage and mobile-element modules that transcend single-locus emm classification.

### Phylons recapitulate established GAS population structure

To determine whether phylon structure corresponded to established GAS classification systems, we compared phylon assignments multilocus sequence typing (MLST), *emm* typing, and *emm*-pattern classification ^14,15,53^. Among the seven lineage-specific taxonomic phylons, concordance with both MLST and *emm* typing was high, with adjusted Rand index (ARI) values of 0.92 and 0.93 and normalized mutual information (NMI) values of 0.89 and 0.90, respectively ^31^. Per-phylon purity confirms that each lineage-specific phylon is dominated by a single sequence type (88–99%) and *emm* type (89–100%), with the *emm3/6/5* clade phylon spanning three related emm types by design (81% combined purity, **Table S5**).

Inclusion of the two composite phylons reduced overall concordance metrics (ARI/NMII approximately 0.75 vs. both MLST and *emm* typing), consistent with their aggregation of numerous low frequency lineages rather than correspondence to a single *emm* or sequence type. However, these composite phylons aligned strongly with higher order *emm*-pattern architecture. Specifically, 94% of strains within the *emm*-patternD-other phylon mapped to D-family *emm* clusters or the closely related clade Y, whereas 86% of strains within the *emm*-patternE-other phylon mapped to E-family *emm* clusters ^12^. Thus, while lineage-specific phylons recapitulate classical emm and MLST structure, the composite phylons capture broader tissue-tropism-associated regulon architectures not resolved by single-locus typing alone. Per-phylon binary concordance metrics are reported in **Table S5**.

Together, these findings demonstrate that NMF recovers biologically meaningful GAS population structure across multiple hierarchical levels, spanning clonal lineage identity, *emm*-pattern organization, and mobile-element-associated accessory genome variation.

### The Sda1 prophage defines the primary split in the GAS accessory genome

To investigate higher-order relationships among phylons, we performed hierarchical exclusive-gene analysis on the binarized L matrix, identifying genes uniquely associated with each branch of the phylon dendrogram (**Fig. 4**). Unexpectedly, the deepest division among taxonomic phylons did not follow the classical emm-pattern framework (A–C versus D versus E). Instead, it separated three phylons, ST28-*emm1*, ST36, and ST63-*emm77,* from all remaining lineages through a shared 27-gene module absent from every other taxonomic phylon. This single split partitions 58% of the taxonomic accessory genome (538 of 924 genes), with 235 genes exclusive to the three-phylon group and 303 genes exclusive to the remaining six phylons. Functional annotation revealed that the shared 27-gene block corresponds to a complete temperate prophage encoding the DNase Sda1 together with integration ^3^), replication ^3^, structural ^3^, lysis ^4^, paratox cassette ^2^, and accessory cargo ^6^ including Sda1 itself, hyaluronidase, hyaluronoglucosaminidase, a secreted protein, glycerate kinase, and nucleoside diphosphate kinase (**Fig. 5; Table S7**).

**Figure 4:**
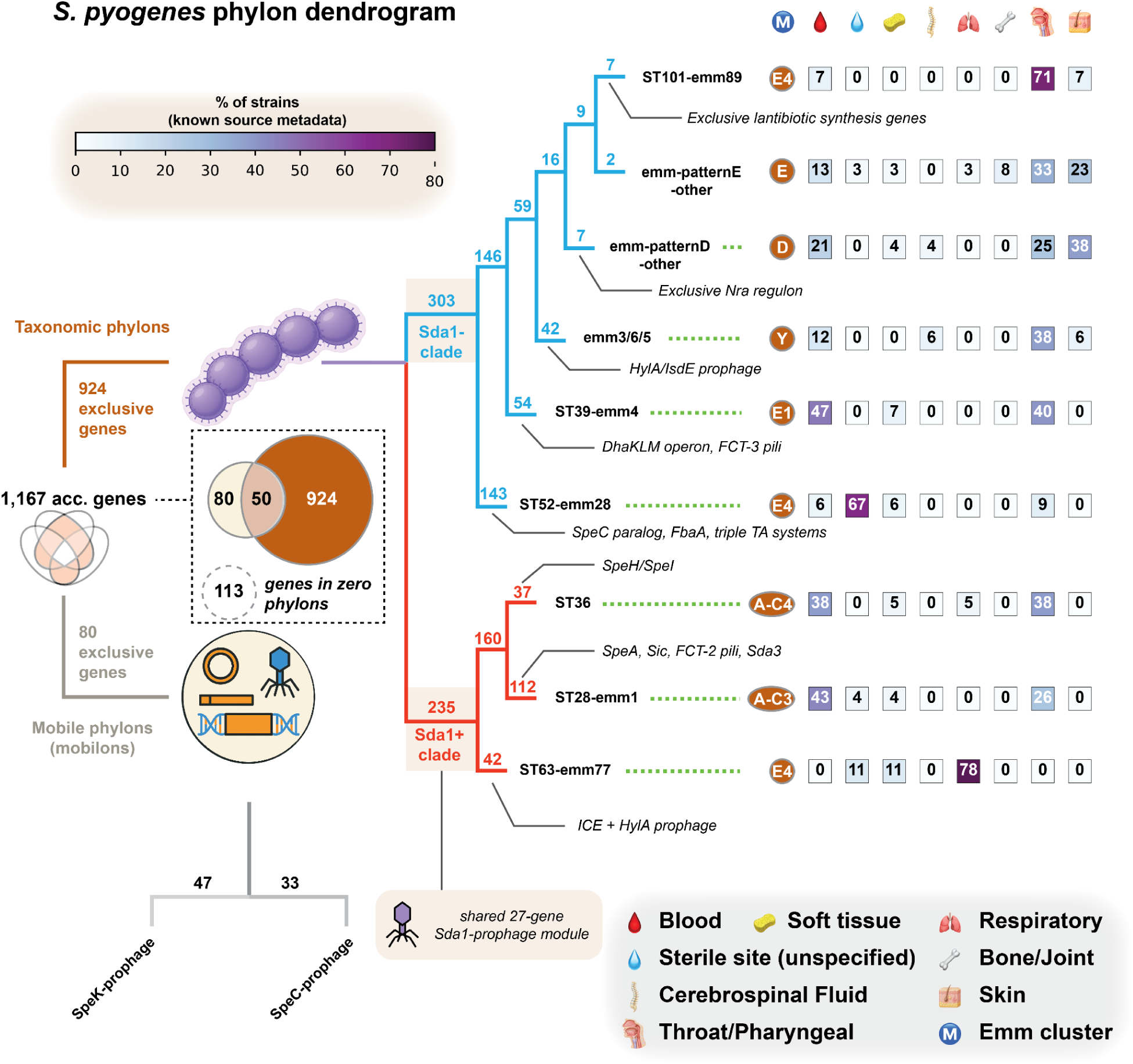
Hierarchical organization of GAS phylons reveals a major Sda1-associated division in the accessory genome. The dendrogram depicts relationships among 11 GAS phylons derived from hierarchical exclusive-gene analysis of the binarized L matrix. Phylons partition into nine taxonomic phylons (orange; 924 exclusive genes) and two mobile phylons, or mobilons (gray; 80 exclusive genes), with 50 genes shared between the two categories. Of the 1,167 accessory genes, 113 are not assigned to any phylon. Branch-adjacent numbers indicate the number of strains exclusively assigned to each phylon. The deepest split among taxonomic phylons separates an Sda1-associated clade comprising ST28-emm1, ST36, and ST63-emm77 from all remaining lineages, bridging the classical A–C/E emm-pattern divide. Right-side annotations summarize representative biological features of each phylon, including emm cluster/pattern assignment and isolation-source distribution (blood, sterile site, soft tissue, cerebrospinal fluid, respiratory system, bone/joint, throat/pharyngeal, and skin). Composite phylons corresponding to emm-pattern D and E architectures aggregate multiple rare lineages sharing common tissue-tropism regulons. Mobilons capture prophage-associated gene modules distributed across multiple taxonomic backgrounds independent of clonal lineage structure.

**Figure 5:**
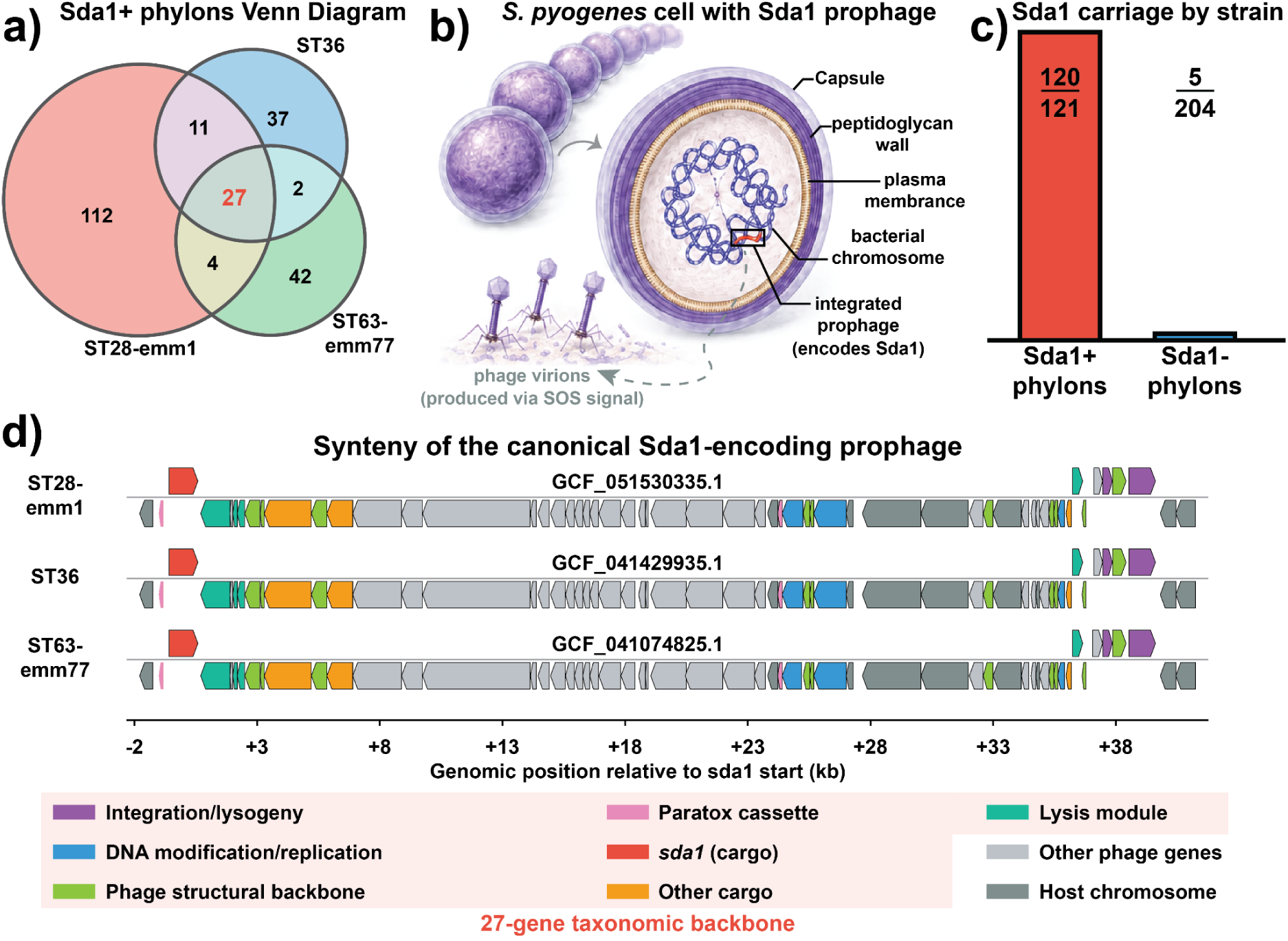
A conserved Sda1-encoding prophage links three major GAS lineages across emm-pattern boundaries. (**a)** Venn diagram showing the distribution of 235 genes exclusive to the three Sda1+ phylons across ST28-emm1, ST36, and ST63-emm77. The shared 27-gene intersection constitutes the conserved Sda1 prophage core, whereas non-overlapping regions represent lineage-specific accessory repertoires, including SpeA, Sic, and FCT-2 pili in ST28-emm1; SpeH/SpeI and PrtF1 in ST36; and a second HylA/IsdE prophage together with an integrative conjugative element (ICE) in ST63-emm77. **(b)** Schematic of a GAS chromosome carrying an integrated Sda1-encoding prophage. Sda1/SdaD2 is a divalent-cation-dependent extracellular DNase that promotes immune evasion through degradation of neutrophil extracellular traps (NETs). **(c)** Per-phylon distribution of sda1 across the 399-genome dataset. Sda1+ phylons (red) carry sda1 at near-fixation (120/121 strains, 99.2%), whereas carriage among all remaining taxonomic phylons is rare (5/204 strains, 2.5%). Mobilons are excluded to avoid double-counting strains assigned to both a taxonomic phylon and a mobilon. **(d)** Cross-phylon synteny of representative Sda1 prophages from ST28-emm1 (GCF_051530335.1), ST36 (GCF_041429935.1), and ST63-emm77 (GCF_041074825.1), selected by highest A-matrix affinity within each phylon. Coordinates are plotted relative to the sda1 start codon (0 kb); genes are colored by functional module. Twenty-five of the 27 shared prophage-core genes are co-localized within a conserved ∼41 kb prophage with preserved geneorder across all three lineages; the remaining two genes (groups_343 and groups_3323) are located elsewhere in the chromosome and are not shown. In all three genomes, the prophage is integrated adjacent to trmB with identical downstream chromosomal organization (trmB–ccrZ–ecsB), consistent with long-term vertical maintenance following ancestral integration.

The 27 genes were tightly co-segregating across the three Sda1+ phylons, with median carriage of 99.2% (range 97.5%–100%) among Sda1+ strains compared with median 4.0% (range 2.5%–22.3%) among non-Sda1+ strains (BH-corrected Fisher’s exact q < 5.2×10^-56^ for all genes; median q = 1.7×10^-87^; **Table S5**). Cross-phylon synteny analysis of representative Sda1+ genomes from each lineage demonstrated that 25 of the 27 genes are co-localized within a conserved ∼41 kb prophage element integrated adjacent to *trmB*, with preserved gene order and modular architecture across all three lineages (**Fig. 5**). The downstream chromosomal flank (*trmB*–*ccrZ*–*ecsB*) was identical in all representatives, consistent with long-term vertical maintenance following ancestral integration ^16,32^. One ST28-emm1 genome additionally carried tandemly integrated Sda1-encoding prophages with near-identical sequence content, consistent with a recent intra-genomic duplication event (**Fig. S6; Table S8**).

Sda1 is a key virulence factor of the hypervirulent *emm1*/M1T1 pandemic clone, where it promotes immune evasion through degradation of the DNA scaffold of neutrophil extracellular traps (NETs) ^21,22^, and has also been identified in *emm12*/ST36 strains ^54,55^. In contrast, carriage within *emm77*/ST63 has not previously been reported. Across the 399-genome dataset, the PanTA *sda1* cluster was carried at near fixation in all three Sda1+ pylons—ST28-emm1 (88/88), ST36 (23/24) and ST63-emm77 (9/9)—whereas carriage in all remaining taxonomic pylons was rare and typically fragmentary. Overall, 120 of 133 total sda1 carriers (90%) belonged to one of the three Sda1+ phylons. Protein-level comparisons further demonstrated near-complete sequence conservation across the shared prophage backbone, with most Sda1 orthologs showing 100% amino-acid identity. These data support a model in which Sda1 carriage is strongly associated with a single conserved prophage module vertically maintained across three otherwise distinct taxonomic lineages.

The Sda1 prophage therefore emerges as a major organizing element of the GAS accessory genome, linking *emm1* (*emm* cluster A–C3), ST36 (*emm* cluster A–C4), and *emm77* (*emm* cluster E4) ^12^ nto a shared Sda1+ clade that transcends the classical A–C/E emm-pattern divide. Despite their shared prophage backbone, each lineage carries a distinct accessory virulence repertoire (**Table 1**), including SpeA/Sic/FCT-2 pili in ST28-*emm1*, SpeH/SpeI/PrtF1 in ST36, and a second HylA/IsdE prophage together with a resistance-associated ICE in ST63-*emm77*. ST63-*emm77* has previously been characterized as a globally circulating pharyngeal lineage enriched for macrolide and tetracycline resistance determinants ^56,57^; its placement within the Sda1+ clade extends the distribution of this NET-evasion module beyond classically invasive lineages and highlights a latent genomic capacity for invasion within a predominantly carriage-associated background.

**Table 1.** Virulence, mobile-element, and ecological characteristics of the seven lineage-specific GAS phylons. Sda1 indicates carriage of the conserved 27-gene Sda1-encoding prophage identified by hierarchical exclusive-gene analysis. “Other superantigens,” “DNases (other),” “Adhesins/pili,” “Immune evasion,” “Other mobile elements,” and “Metabolic/competition” summarize the principal lineage-associated accessory gene content derived from the binarized L matrix (see **Methods**); mobilon-associated genes are indicated in parentheses. Composite phylons (emm-patternD-other and emm-patternE-other) are excluded because they aggregate multiple rare lineages and are analyzed separately in the regulon-recovery section.

| Phylon | ST28-emm1 | ST36 | ST63-emm77 | ST52-emm28 | ST39-emm4 | emm<br>3/6/5 | ST101-emm89 |
| --- | --- | --- | --- | --- | --- | --- | --- |
| # strains | 88 | 24 | 9 | 33 | 28 | 31 | 18 |
| # excl. genes | 112 | 37 | 42 | 143 | 54 | 42 | 7 |
| Sda1 | + | + | + | - | - | - | - |
| Other<br>Superantigens | SpeA | SpeH,<br>SpeI | - | SpeC<br>paralog<br>(lineage-specific) | SpeC<br>(mobilon) | - | - |
| DNases (other) | Sda3 | - | - | - | - | - | - |
| Adhesions/<br>pili | FCT-2<br>pilus,<br>Scl1,<br>Scl2,<br>FbpA | PrtF1/Sfb<br>I, distinct<br>pilus<br>locus | - | FbaA | FCT-3 pili | - | - |
| **Immune evasion** | Sic | - | - | - | - | - | - |
| **Other MGEs** | - | - | 2nd prophage (HylA, IsdE); ICE | - | SpeC prophage (mobilon) | HylA/IsdE prophage | - |
| **Metabolic/competition** | - | - | - | 3 TA systems (MazF, RelE/RelB, HicB) | DhaKLM (glycerol/DHA operon) | - | LanM lantibiotic system |
| **Dominant isolation site** | Blood (45%) | Throat/P haryngeal (42%) | Throat/P haryngeal (78%) | Sterile site (67%) | Blood (43%) | Throat/P haryngeal (46%) | Throat/P haryngeal (75%) |
| **Invasive isolation rate** | 73% | 53% | 22% | 90% | 57% | 46% | 17% |
| **Invasive (n/N)** | 16/22 | 10/19 | 2/9 | 27/30 | 8/14 | 6/13 | 2/12 |

### Two independent mobilons encode superantigen prophages, with recurrent acquisition of divergent *speC*/*mf2* cassettes

The two mobilons identified by NMF correspond to phylogenetically distinct prophage elements encoding different superantigen repertoires. Of the 80 mobilon-exclusive genes, 47 belong uniquely to the SpeK-associated mobilon and 33 to the SpeC-associated mobilon, with no genes shared between them (**Fig. 4**). This absence of gene-level homology indicates unrelated phage backbones: the SpeK prophage belongs to the TP901-1/PBSX-terminase family (Φ315.4-like), whereas the SpeC prophage carries P27-family terminase and HK97-gp10 morphogenesis machinery ^16,58^.

The SpeK-prophage mobilon exhibits a restricted lineage distribution, with 11 of 13 carriers (85%) occurring within the emm3/6/5 phylon, and encodes superantigen SpeK together with the phospholipase Sla ^2,58^. In contrast, the SpeC mobilon spans 16 *emm* types distributed across five taxonomic phylons, with strongest enrichment in ST39-*emm4* (15 of 29 carriers, 52%). Within this lineage, SpeC and the DNase Mf2 approach fixation, consistent with stable vertical maintenance of the prophage following acquisition. No genome carried both mobilons simultaneously, suggesting functional redundancy or superinfection exclusion between these prophage systems ^33^.

Beyond the mobilon-associated copies, the pangenome revealed that SpeC circulates as three sequence-divergent paralog families carried on independent prophage backbones. PanTA separated this into speC_01049 (n = 90 genomes, carried by the SpeC-prophage mobilon and at fixation in ST39-*emm4*), speC (n = 49 genomes, at fixation in ST52-*emm28* with additional carriage in ST63-*emm77* and *emm*-pattern E lineages), and *speC_01050* (n = 44 genomes, distributed across *emm3/6/5*, *emm*-patternE-other, and *emm*-patternD-other). Each was paired with a corresponding DNase mf2 paralog that co-occurred without discordance across all carrier genomes (mf2_01051: n = 90; mf2: n = 49; mf2_01052: n = 44). The three paralog pairs differed by >20% amino-acid sequence divergence yet retained identical genomic organization: in representative genomes, *speC* and *mf2* occurred in conserved head-to-tail orientation ^18^ immediately downstream of phage structural and lysis modules and adjacent to a small Paratox gene (**Fig. 6**) Despite extensive divergence among the surrounding prophage backbones, all three prophages independently encoded hyaluronidase (HylA) upstream of the cargo cassette and integrated at distinct chromosomal attachment sites, indicating multiple independent acquisition events (**Table S9**).

**Figure 6.**
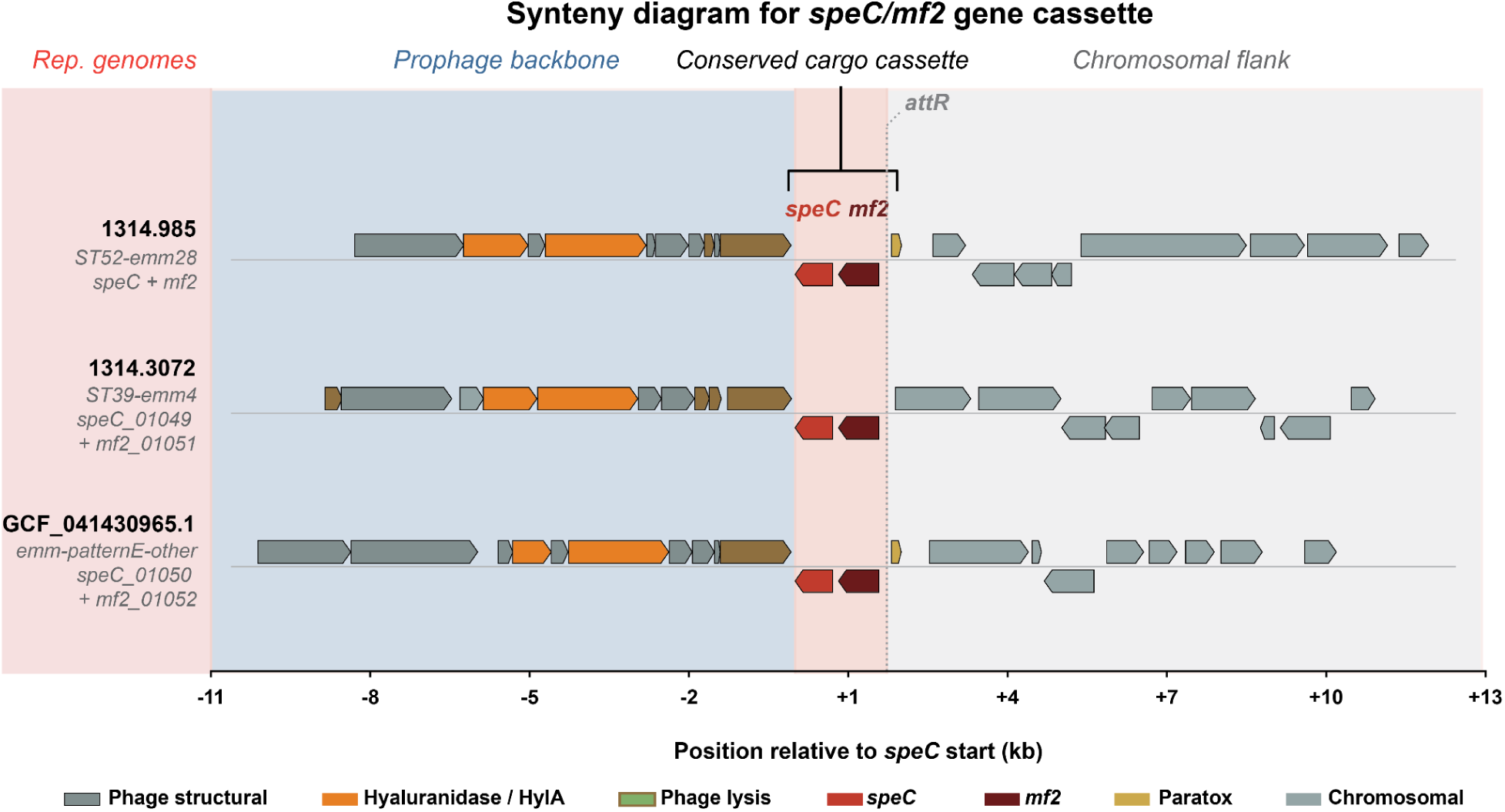
Recurrent acquisition of divergent speC/mf2 virulence cassettes across unrelated prophages. Synteny of the speC/mf2 region in representative genomes from the three paralog-defined carrier groups identified by PanTA clustering: 1314.985 (ST52-emm28; speC + mf2), 1314.3072 (ST39-emm4; speC_01049 + mf2_01051), and GCF_041430965.1 (emm-patternE-other; speC_01050 + mf2_01052). Coordinates are plotted relative to the speC start codon (0 kb). Genes are colored by functional category. In all three prophages, speC and mf2 occur in conserved head-to-tail orientation immediately downstream of phage structural and lysis genes and adjacent to a small Paratox gene. Despite major divergence among prophage backbones, all three independently encode hyaluronidase (HylA) upstream of the toxin cassette, consistent with convergent selection of a conserved virulence module. The prophages integrate at distinct chromosomal loci, indicating independent acquisition events. The GCF_041430965.1 prophage is integrated in reverse orientation relative to the other two genomes and was reverse-complemented for visualization. Per-gene locus tags, genomic coordinates, and functional annotations are provided in Table S8.

Together, these findings support convergent evolutionary selection of a conserved speC/mf2/HylA virulence cassette across unrelated prophage lineages. Although carried on structurally distinct prophages integrated at different chromosomal loci, the repeated preservation of this toxin-DNase-hyaluronidase module suggests that coordinated superantigen activity, extracellular DNase function, and tissue-spreading capacity provide a selectively advantageous combination within diverse GAS genetic backgrounds ^17^.

### Distinct GAS lineages encode alternative virulence architectures associated with invasion and niche adaptation

Hierarchical exclusive-gene analysis revealed that each taxonomic phylon carries a unique combination of superantigens, adhesins, immune evasion factors, and metabolic functions (**Table 1**). Although the Sda1+ phylons share a common NET evasion prophage, their broader virulence repertoires differ substantially. ST28-emm1 carries the canonical M1T1-associated complement of SpeA, Sic, FCT-2 pili, and the collagen-like adhesins Scl1/Scl2 ^2,20^. ST36 instead encodes the alternative superantigen pair SpeH/SpeI together with the adhesin PrtF1 on a distinct prophage backbone. ST63-*emm77* lacks superantigens among its exclusive genes, but carries a second prophage encoding HylA and IsdE together with an integrative conjugate element (ICE), consistent with a genomic architecture emphasizing tissue penetration, iron acquisition, and antimicrobial resistance atop the shared Sda1-mediated NET-evasion platform ^2,56^.

Among Sda1-negative phylons, ST52-*emm28* exhibited the highest invasive isolation frequency in the dataset (90% invasive source sites among strains with metadata) despite lacking the Sda1 prophage. Instead, this lineage carries a distinct SpeC paralog together with the fibronectin-binding adhesin FbaA and an unusually dense toxin-antitoxin repertoire, MazF, RelE/RelB, HicB, representing the highest toxin-antitoxin system density observed in any phylon. ST39-*emm4* (57% invasive) combines broadly disseminated SpeC mobilon with FCT-3 pili and a dedicated glycerol/dihydroxyacetone utilization operon (DhaKLM), potentially supporting enhanced fitness in the pharyngeal niche ^59^. The *emm3/6/5* clade phylon, by contrast, is dominated by a structurally distinct prophage encoding HylA and IsdE, representing convergent evolution of tissue-spreading and iron-acquisition functions on independent phage backbones ^17^.

At the opposite end of the invasive spectrum, ST101-*emm89* was predominantly associated with throat isolates (75%) and carried a lineage-specific lantibiotic biosynthesis system (LanM) ^49,50^, consistent with a niche-competition strategy based on bacteriocin production rather than invasive virulence.

Together, these findings demonstrate that invasive potential in GAS is not linked to a single virulence determinant or prophage lineage, but instead emerges through multiple lineage-specific virulence architectures combining distinct toxins, adhesins, immune-evasion systems, metabolic programs, and mobile genetic elements. Notably, some lineages with substantial genomic capacity for invasion, including ST63-*emm77*, remain predominantly associated with pharyngeal carriage, suggesting that latent invasive potential may circulate within clinically underappreciated genomic backgrounds.

### NMF recovers emm-pattern regulons and recombination-driven emm-switching

Two of the eleven phylons, *emm*-patternD-other and *emm*-patternE-other, did not correspond to single dominant *emm* types but instead grouped diverse lineages sharing common regulatory architectures associated with tissue tropism. The *emm*-patternD-other composite was enriched for skin-associated specialist lineages characterized by the Nra transcriptional regulator, whereas *emm*-patternE-other grouped generalist lineages associated with the RofA regulon and mixed throat/skin tropism ^14^.

Despite their polyclonal composition, these composite phylons recovered coherent biological modules. The emm-patternD-other phylon was defined by seven exclusive genes constituting the canonical Nra regulon, ^14^ including Nra itself, which replaces Mga in the pattern D arrangement, together with Enn ^13^, SfbII/PrtF2, SilD, IFS, and two additional regulatory proteins. In contrast, *emm*-patternE-other retained only two exclusive genes after hierarchical filtering, with its identity driven primarily by the broader Mga/Mrp/SfbI regulon architecture shared across generalist emm-pattern E strains. Concordance with emm-cluster classification supported this partitioning: 94% of *emm*-patternD-other strains mapped to D-family emm clusters (D1–D5) or the related clade Y, whereas 86% of *emm*-patternE-other strains mapped to E-family clusters (E1–E6) ^12^. These findings demonstrate that NMF can recover biologically coherent tissue-tropism regulons directly from accessory genome structure without prior annotation.

The decomposition additionally detected a recombination-driven emm-switching event within the ST36 phylon. This phylon grouped 18 *emm12* strains (17 ST36 and one ST242) together with five *emm82*/ST36 strains into a single co-occurring gene set, indicating conservation of the underlying chromosomal backbone despite divergence at the *emm* locus. One additional ST36 isolate could not be assigned by emmtyper and was excluded from the *emm12*/*emm82* comparison. Notably, this grouping emerged entirely through unsupervised decomposition without prior information regarding *emm* switching. Recent genomic surveillance studies independently identified *emm82*/ST36 isolates arising through recombinational replacement of the *emm12 emm*-region ^60^, supporting the ability of the phylon framework to detect lineage-level recombination events directly from pangenome structure.

## Discussion

The decomposition of the GAS accessory genome into 11 phylons reveals a major axis of genomic organization not captured by conventional typing schemes. For more than three decades, *emm* typing has served as the foundation of GAS epidemiology, classifying strains according to sequence variation at a single surface-antigen locus. Our results demonstrate that the deepest partition in the accessory genome is instead defined by prophage content: the presence or absence of a conserved 27-gene Sda1-encoding prophage supersedes *emm* type and *emm*-pattern classification as the primary organizer of accessory-genome structure. This does not diminish the epidemiologic utility of *emm* typing; the strong concordance between lineage-specific phylons and *emm* types (ARI = 0.93) confirms that *emm* typing captures authentic clonal structure. Rather, it reveals an additional layer of genomic organization in which shared mobile genetic elements unite lineages that appear unrelated by single-locus classification alone.

The Sda1 prophage appears to represent a remarkably stable lysogenic association between prophage and host. Multiple lines of evidence support long-term vertical maintenance rather than repeated independent acquisition: the prophage occupies a conserved *attB* site adjacent to *trmB* in all three Sda1+ lineages, the 27-gene module maintains identical gene order across phylogenetically distinct backgrounds, and the *sda1* cargo gene shows complete protein conservation across 120 of 121 Sda1+ strains. The tandemly duplicated prophage observed in strain 1314.4589 further underscores the structural stability of the element. This persistence is biologically plausible given the established role of Sda1 in extracellular degradation of neutrophil extracellular traps (NETs), providing a continuous immune-evasion advantage during colonization and invasive infection. Together, these observations support a model in which a highly conserved prophage module has become stably embedded within multiple successful GAS lineages.

This framework connects directly to the covRS selection model ^22^: Sda1-mediated NET evasion creates a selective bottleneck favoring covRS-mutant subclones with upregulated capsule and superantigen expression, driving the transition from colonization to invasive disease. If emm77/ST63 carries Sda1 at fixation alongside HylA, IsdE, and an antimicrobial-resistance ICE, the covRS selection model predicts that emm77 populations are subject to the same selective pressure for covRS mutation that drove the M1T1 pandemic expansion, yet emm77 remains predominantly pharyngeal in clinical surveillance (78% throat isolates, 22% invasive). This gap between latent genomic capacity and realized clinical phenotype suggests that additional factors beyond the Sda1-covRS axis (host immune status, co-colonization dynamics, or as-yet-unidentified regulatory constraints) modulate the transition to invasive disease. Surveillance strategies that directly track Sda1 carriage across circulating GAS populations, rather than relying solely on emm type as a surrogate for invasive potential, may therefore provide earlier insight into lineages poised for invasive emergence.

The discovery of three sequence-divergent SpeC paralogs on independent prophage backbones holds important implications for molecular diagnostics and vaccine development. PCR-based detection assays and antigen-based vaccine candidates designed against one SpeC variant may fail to detect or protect against the others. More fundamentally, the strict co-occurrence of *speC* and *mf2* across three paralog pairs at <80% amino-acid identity, combined with their conserved HylA association on structurally distinct prophages integrated at three different chromosomal sites, demonstrates that superantigen-encoding capacity is a convergent cargo strategy adopted independently by multiple phage lineages, not a marker of shared phage ancestry. This convergence extends to the chromosomal level: the *emm3/6/5* and *emm77* prophages independently acquired hyaluronate lyase and IsdE on structurally unrelated backbones, suggesting that certain virulence gene combinations are strongly selected regardless of the vehicle that delivers them.

Comparison of lineage-specific accessory genomes revealed that invasive potential in GAS is achieved through multiple distinct virulence architectures rather than through a single conserved pathogenic program. Although the three Sda1+ phylons share a common NET-evasion platform, they diverge extensively in their superantigens, adhesins, immune-evasion factors, and mobile genetic elements. Conversely, ST52-*emm28* achieves the highest invasive isolation rate in the dataset despite lacking Sda1, instead combining a distinct SpeC paralog, FbaA-mediated adhesion, and an unusually dense toxin-antitoxin repertoire. Together, these findings suggest that invasive disease in GAS represents a convergent phenotype emerging from multiple independent genomic solutions rather than from a universal virulence determinant.

A distinctive feature of GAS genome evolution is that prophage cargo contributes simultaneously to clonal lineage identity and mobile genetic exchange — unlike species such as *E. coli* or *S. aureus*, where plasmids and genomic islands largely occupy separable evolutionary niches and can be distinguished from chromosomal signal by replicon or compositional features alone ^31^. In GAS, the same category of genes — prophage-encoded virulence determinants — defines both stable lineage structure and ongoing horizontal transfer, creating superimposed inheritance signals within a single chromosomal backbone. That phylon decomposition recovered lineage structure, mobile prophage elements, an independently validated *emm*-switching event, and tissue-tropism regulons from binary gene presence/absence data alone, without phylogenetic reconstruction or prior functional annotation, suggests that this superimposition is organized rather than intractable. The additive structure of NMF, in which each genome is reconstructed as a non-negative combination of co-occurring gene sets, appears well matched to the layered architecture by which GAS genomes are assembled through vertical inheritance and prophage acquisition. This property may extend to other highly lysogenized pathogens whose accessory genomes are similarly dominated by integrated mobile elements.

Several limitations should be considered. The dataset of 399 complete genomes underrepresents tropical and skin-tropic lineages, limiting the resolution of the emm-pattern D and E composite phylons; the extraction of ST101-emm89 as an independent phylon at rank 11 directly demonstrates that these composites likely reflects incomplete sampling of globally circulating lineages, and we anticipate that future decompositions with larger genome collections will resolve additional lineage-specific phylons. The restriction to complete genomes and the reliance on emmtyper (which classifies by M protein sequence rather than *mga* locus gene arrangement) impose further constraints. NMF identifies co-occurring gene sets but does not establish causality; associations between gene combinations and invasive disease require experimental validation. Finally, the absence of transcriptomic data precludes confirmation that co-occurring genes are co-regulated; integration with iModulon-based transcriptomic decomposition, connecting evolutionary modularity (phylons) with regulatory modularity (iModulons), would provide a complementary perspective on how the *S. pyogenes* genome is both assembled and deployed.

In summary, the phylon framework resolves GAS population structure simultaneously at the levels of clonal lineage, prophage content, and tissue-tropism regulons from a single decomposition of the accessory genome. The identification of a vertically maintained Sda1 prophage as the dominant axis of accessory-genome organization, together with the discovery of convergent superantigen cargo architectures across unrelated prophages, argues that surveillance efforts should move beyond single-locus typing toward direct tracking of accessory-genome configurations associated with immune evasion and invasive potential. As invasive GAS disease continues to re-emerge globally, understanding how virulence modules circulate, recombine, and stabilize within successful lineages may prove critical for anticipating pathogen emergence and designing durable intervention strategies.

## Materials and Methods

### Experimental design

This study is entirely computational and was designed to test whether the *S. pyogenes* accessory genome possesses an underlying modular organization recoverable from gene presence/absence patterns alone, and whether non-negative matrix factorization (NMF) can disentangle the superimposed signals of clonal descent and prophage-mediated gene flow in a highly lysogenized species. We assembled a curated compendium of 399 high-quality complete *S. pyogenes* genome sequences (Spyo-GENOMiCUS-400) from the BV-BRC and NCBI RefSeq databases, applied NMF to the binary accessory-gene presence/absence matrix to identify co-occurring gene sets (phylons), and characterized the resulting structure through three prespecified analyses: (i) concordance against established classification systems (MLST sequence types, emmtyper-derived *emm* types, and emm-pattern classifications), (ii) hierarchical exclusive-gene analysis to quantitatively partition the accessory genome into vertically and horizontally inherited components, and (iii) targeted validation of the Sda1 prophage distribution by tBLASTn. The NMF rank was prespecified by a two-stage calibration procedure (Stage 1: CD-HIT-derived pangenome with optNMF; see Pangenome generation: two-stage approach) before downstream biological interpretation. No genome sequences were generated for this study; all data are publicly available.

### Bacterial genome sequences

The primary dataset (Spyo-GENOMiCUS-400) comprises 399 high-quality, fully circularized *Streptococcus pyogenes* genome sequences downloaded from the BV-BRC database (https://www.bv-brc.org/) and NCBI RefSeq (https://www.ncbi.nlm.nih.gov/refseq/) as of April 2026. BV-BRC yielded 2,867 genome sequences and NCBI contributed 835 complete genome assemblies. Metadata including isolation source, geographic location, collection date, host information, and strain identifiers were retrieved from both databases and merged into a unified metadata table. Geographic location was consolidated by combining the “isolation_country” and “geographic_location” metadata fields, with manual curation of ambiguous entries.

### Gathering and processing of genome sequence data

We downloaded the metadata of all genomes available on BV-BRC and NCBI RefSeq classified as *Streptococcus pyogenes*. Genome sequences labeled “Complete” were filtered by their assembly metrics: L50 score required to equal 1 and N50 score required to exceed 1,500,000. CheckM (v1.2.2) contamination (<3%) and completeness (>97%) scores were applied to filter genomes further. The resulting collection was deduplicated and collated for further quality control via Mash filtration (described below). Ultimately, only complete (fully circularized) genome sequences were retained for pangenome analysis to ensure the highest quality gene presence/absence matrix **P**.

Following initial quality control, 402 complete genome sequences remained. An additional 3 strains were excluded based on anomalous gene composition: strains with core, accessory, or rare gene counts exceeding 4 standard deviations from the median in the core-accessory-rare (CAR) genome breakdown were removed as statistical outliers. The final curated dataset comprises 399 high-quality complete *S. pyogenes* genome sequences, designated Spyo-GENOMiCUS-400.

### Genome annotation with Bakta

All 402 genomes (pre-outlier removal) were annotated using Bakta (v1.12.0) (28) with the full database (v6.0). Bakta was run with the following parameters: “--complete”, “--gram +”, “--genus Streptococcus”, “--species pyogenes”, “--translation-table 11”, “--verbose”, with 8 threads per genome. The “--gram +” flag enabled DeepSig-based signal peptide predictions appropriate for this Gram-positive species. Annotation was parallelized across genomes using GNU Parallel (8 concurrent jobs × 8 threads) on a 128-core server. All annotated genomes were screened by the *S. pyogenes* PubMLST schema through the mlst GitHub package to identify sequence types for all strains.

### Mash filtration and analysis

All downloaded genomes were compared to generate pairwise Mash distance values using Mash (v2.3) (29) with a sketch size of 10,000. A Mash distance threshold of 0.05 (corresponding to a 70% DNA-DNA reassociation value and the historical cutoff between bacterial species) was used to identify and remove any genomes falling outside the *S. pyogenes* species boundary. Mash distance values were then converted to Pearson correlation coefficients, which were subsequently converted to Pearson correlation distances for hierarchical clustering using Ward’s minimum variance method. The number of clusters formed at the optimal threshold was saved as a potential candidate rank for NMF decomposition. This Mash-based clustering also informed initial metadata exploration, including the identification of 12 filtered Mash clusters that were subsequently compared against phylon assignments.

### Pangenome generation: two-stage approach

We used a two-stage pangenome construction workflow that separates parameter estimation from final biological analysis. In Stage 1, a CD-HIT-derived pangenome was used to estimate the core/accessory/rare genome boundaries and to calibrate the NMF rank. In Stage 2, the final pangenome was generated with PanTA, and all downstream analyses were performed on the PanTA-derived gene families at the calibrated rank. This design avoids circularity: the gene-family definitions used for final biological interpretation do not themselves determine the pangenome boundaries or the decomposition rank.

### Stage 1: CD-HIT-based pangenome for boundary determination and rank calibration

Bakta-annotated genomes were initially clustered into gene families using CD-HIT (v4.8.1), with sequence similarity and alignment coverage cutoffs of 80%, consistent with standard pangenome practice ^31,36^. This sequence-similarity-based pangenome served two purposes. First, its gene-frequency distribution was used to fit the double-exponential model of Hyun et al. ^36^ for determining core, accessory, and rare genome boundaries. Second, the resulting accessory-genome presence/absence matrix was used as input for optNMF rank calibration.

This separation is necessary because synteny-aware pangenome tools improve biological gene-family resolution but alter the frequency distribution used for boundary and rank estimation. PanTA distinguishes true orthologs from positionally distinct paralogs and separates mobile-element insertions by genomic context. This correctly prevents transposable elements and recent paralogs from being collapsed into artificially high-frequency gene families, but it also shifts the inflection points of the cumulative gene-frequency distribution. Similarly, synteny-aware clustering compresses the effective dimensionality of the accessory genome, causing optNMF applied directly to the PanTA matrix to select a lower reconstruction optimum that captures major taxonomic structure but may compress mobilons into broader components.

### Rank calibration and decomposition stability

We calibrated NMF rank using the optNMF algorithm applied to the CD-HIT-derived accessory-genome matrix. Candidate ranks were evaluated by performing NMF, binarizing the resulting L and A matrices, reconstructing the binary accessory-genome matrix, and comparing the reconstruction to the original binary matrix. For each rank, reconstruction performance was summarized using precision, recall, F1-score, Matthews correlation coefficient (MCC), false-positive rate, and reconstruction Jaccard.

The reconstruction Jaccard was computed entrywise between the original binary accessory-gene matrix **P**_orig and the binarized reconstruction **P**_reconstr:

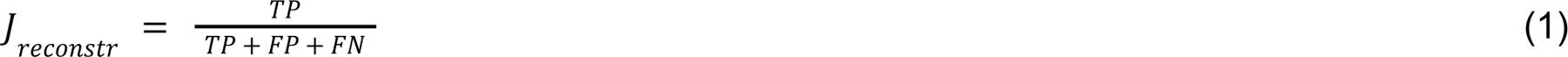

where TP denotes the number of true positives (both entries in **P**_orig and **P**_reconstr are 1), FP denotes the number of false positives (the entry in **P**_orig is 0 but the entry in **P**_reconstr is 1), and FN denotes the number of false negatives (the entry in **P**_orig is 1 but the entry in **P**_reconstr is 0).

The AIC proxy used for optNMF rank selection was computed from the reconstruction Jaccard distance and a rank-dependent parameter penalty:

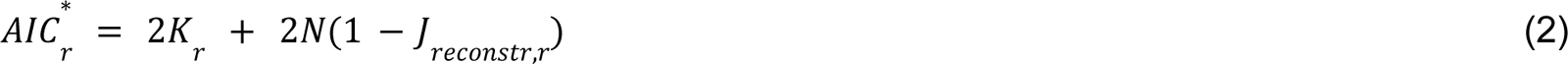

where r is the NMF rank, N is the total number of entries in the accessory-gene matrix, and K_r = 2r(m + n) is the conservative parameter count for the two NMF factor matrices for a matrix with m genes and n genomes. Because this is an AIC proxy rather than a likelihood-derived AIC, values were interpreted only within a given pangenome matrix and rank sweep, not as absolute model likelihoods across different gene-family definitions.

For Spyo-GENOMiCUS-400, optNMF applied to the CD-HIT-derived accessory-genome matrix selected rank 11.

### Stage 2: PanTA-based pangenome for analysis

The final pangenome was generated using PanTA (https://github.com/amromics/panta) with the following parameters: “-i 0.8 --AL 0.8 -b diamond -t 64”, corresponding to a minimum percent identity of 80% and a minimum alignment length coverage of 80% for gene clustering, using DIAMOND as the alignment engine. PanTA performs ortholog/paralog splitting based on synteny and genomic context, which is particularly important for resolving gene families that share high sequence similarity but occupy distinct genomic positions or have different evolutionary histories: a critical consideration in *S. pyogenes*, where multiple prophages carry structurally similar but functionally distinct virulence genes (e.g., three SpeC paralogs on independent prophage backbones).

The core, accessory, and rare genome boundaries determined from the CD-HIT pangenome in Stage 1 were applied to the PanTA-derived gene-frequency distribution. The resulting PanTA accessory-genome matrix was then decomposed by NMF at the CD-HIT-calibrated rank of 11. All downstream analyses, including phylon characterization, functional annotation, hierarchical exclusive-gene analysis, and concordance assessment, were performed on the PanTA-derived pangenome.

### Defining the core, accessory, and rare genomes

The core, accessory, and rare genomes were defined using the cumulative gene distribution plot, following methods established by Hyun et al. (2). Briefly, the gene frequency distribution (number of strains containing each gene) forms a characteristic S-shaped cumulative curve with identifiable inflection points. This curve was fitted to a double-exponential model (median absolute error = 43.41).

The core genome boundary was defined by taking the highest endpoint of the cumulative distribution and traveling 90% of the distance from the inflection point to this endpoint, corresponding to the elbow in the plot. This yielded a core genome threshold of ≥98.2% strain presence (≥391/399 genomes), identifying 1,292 core genes. The rare genome boundary was defined analogously using the lowest endpoint, yielding a threshold of <8.2% strain presence (<33 genomes), identifying 6,405 rare genes. The accessory genome comprises genes between these two thresholds: those found in 8.2–98.2% of all strains (33–391 genomes), totaling 1,167 genes. The total pangenome comprises 8,864 gene families.

### Non-negative matrix factorization

The accessory genome matrix **P** (1,167 genes × 399 strains) was decomposed using non-negative matrix factorization (NMF). The scikit-learn implementation of NMF (v1.x) was used with the following parameters: rank of 11, Frobenius norm objective, coordinate descent solver, initialization of “nndsvd,” and a maximum iteration limit of 10,000. NNDSVD initialization is deterministic: given the same input matrix, it produces identical factor matrices on every run, eliminating the seed-dependent variability that motivates consensus NMF approaches.

### Normalization and binarization

#### Normalization

For each column in **L**, the 99th percentile value was calculated, and every value in that column was divided by this value, ensuring all but a few values fell between 0 and 1. To maintain reconstruction consistency, the corresponding rows in the **A** matrix were multiplied by the same normalization factors. This normalization ensures that the normalized matrix **L**’, when multiplied by the normalized matrix **A**’, results in the same reconstructed **P** matrix as that of the product of the original **L** and **A** matrices.

#### Binarization

The normalized **L** and **A** matrices were binarized using k-means clustering (k = 3), implemented in scikit-learn. For each column of **L**, gene weightings were segregated into three clusters; genes in the cluster with the highest mean were set to 1 (indicating membership in the phylon), while genes in the other two clusters were set to 0. The same procedure was applied to the rows of the **A** matrix. The choice of k = 3 provides a conservative binarization: the three clusters separate strong positive signals from intermediate values and near-zero background, assigning membership only to the highest cluster. This conservatism is appropriate given the bimodal distribution of continuous weights in both **L** and **A**, which converge toward values near unity or near zero during optimization, leaving few ambiguous intermediate values.

#### Phylon characterization

Phylons were characterized and named using the binarized **A** matrix. Strains with “high affinity” for a phylon were defined as those with a binarized entry of 1 for that phylon. Phylons were first classified as taxonomic or mobile based on strain co-occurrence patterns: phylons whose high-affinity strains never overlapped with those of another phylon were classified as taxonomic; phylons whose high-affinity strains systematically co-occurred with a taxonomic phylon were classified as mobile (mobilons). Taxonomic phylons were named based on the predominant MLST sequence type and emmtyper-derived *emm* type of their constituent strains. Composite phylons aggregating multiple rare lineages were named based on their predominant emm-pattern classification (emm-patternD-other, emm-patternE-other). Mobilons were named based on their dominant virulence cargo gene (SpeK-prophage, SpeC-prophage).

#### *emm* typing

*In silico emm* typing was performed on all 399 genome assemblies using emmtyper (v0.2.2; https://github.com/MDU-PHL/emmtyper), installed from the GitHub repository. emmtyper was run with default parameters using the BLAST workflow against the CDC *emm* subtype database bundled with the tool. emmtyper reports the predicted *emm* type, possible *emm*-like alleles (enn, mrp), and the EMM cluster assignment for each genome. emm-pattern assignments (A-C, D, E) were inferred from the emm cluster designations: clusters prefixed with “A-C” were assigned to pattern A-C, clusters prefixed with “D” to pattern D, clusters prefixed with “E” to pattern E, and “Single protein cluster clade Y” entries to clade Y.

#### Hierarchical exclusive-gene analysis

To investigate the relationships among the 11 phylons, we performed a hierarchical exclusive-gene analysis on the binarized **L** matrix. Phylons were first clustered by Ward’s minimum variance method on the **L** matrix columns. At each split of the resulting dendrogram, we identified genes exclusively present in one group (i.e., assigned to ≥1 phylon in the group in the binarized **L** matrix) and absent from all phylons in the other group. This procedure was applied recursively at each level of the hierarchy, with the gene pool at each level restricted to the exclusive genes inherited from the parent split. This constraint ensures that exclusive gene counts at child nodes sum to the parent node count, enabling quantitative accounting of the entire accessory genome through the dendrogram.

#### Sda1 validation by tBLASTn

To validate the identification of Sda1 in emm77/ST63, the Sda1/SdaD2 protein sequence was retrieved from UniProtKB (accession Q675N6, 390 amino acids). All 399 genome assemblies were concatenated into a single FASTA file and formatted as a nucleotide BLAST database using makeblastdb (BLAST+ v2.17.0). tBLASTn was performed with an e-value threshold of 1×10⁻¹⁰ and default parameters. Hits were classified as orthologous if they exhibited ≥80% amino acid identity over ≥80% query coverage (≥312/390 amino acids aligned). Hits below these thresholds were classified as paralogous (e.g., other DNase family members). The distribution of Sda1 orthologs was cross-referenced against phylon assignments from the NMF decomposition and against the PanTA gene presence/absence matrix for the *sda1* gene cluster.

#### Concordance analysis

Concordance between NMF-derived phylon assignments and established typing schemes (MLST, *emm* type, emm pattern) was quantified using the adjusted Rand index (ARI) and normalized mutual information (NMI), computed using scikit-learn. For concordance calculations, only strains assigned to exactly one taxonomic phylon (excluding mobilons) were included. Concordance was computed both for all nine taxonomic phylons (global concordance) and for the seven lineage-specific phylons only (excluding the two composite emm-pattern phylons), to assess the effect of composite phylon aggregation on global concordance metrics. Per-phylon purity was calculated as the fraction of strains within each phylon belonging to the most common ST, *emm* type, or emm-pattern classification. Per-phylon binary concordance was additionally computed for each phylon as a 2 × 2 partition (in-phylon vs not, in-truth-class vs not) using ARI, NMI, sensitivity, and positive predictive value (PPV / purity), with strains lacking the relevant truth label dropped from the denominator (**Tables S5, S6**). For the composite phylons (emm-patternD-other, emm-patternE-other), concordance with the emm-pattern classification is directional: each composite phylon captures the residual pattern-D or pattern-E lineages that are not absorbed by the lineage-specific phylons. Partition-level ARI/NMI between a composite phylon and its parent emm-pattern is therefore expected to be low even when phylon-to-pattern purity is high, and we summarize composite-phylon concordance using purity (the fraction of phylon members assigned to the expected pattern, restricted to strains with an interpretable emmtyper output) rather than partition-level metrics. For the emm3/6/5 clade phylon, which spans three related *emm* types by design, combined purity (emm3 + emm5 + emm6) was reported. For the ST36 phylon, which groups emm12 and emm82 strains sharing the same sequence type, MLST purity alone was reported.

#### Sda1 prophage enrichment analysis

To quantify the association between the 27-gene Sda1 prophage core and Sda1+ phylon membership, we performed a gene-by-gene enrichment analysis. For each of the 27 genes identified by hierarchical exclusive-gene analysis as present in all three Sda1+ phylons and absent from all other phylons, we constructed a 2×2 contingency table comparing gene presence/absence in Sda1+ strains (n = 121) versus non-Sda1+ strains (n = 278). Statistical significance was assessed by one-tailed Fisher’s exact test (alternative = greater), and p-values were corrected for multiple testing using the Benjamini-Hochberg procedure at FDR < 0.05. Log-odds ratios were computed with Haldane-Anscombe correction (0.5 added to all cells) to handle zero counts. Results are reported in Table S7.

#### Prophage synteny analysis

To visualize the conservation of prophage architecture across lineages, we performed synteny analysis on representative genomes selected by highest A-matrix affinity for their respective phylons. For the Sda1-encoding prophage (Fig. 5d), three representative genomes were used: GCF_051530335.1 (ST28-emm1), GCF_041429935.1 (ST36), and GCF_041074825.1 (ST63-emm77). For the speC/mf2 virulence cassette (Fig. 6), three representative genomes were used: 1314.985 (ST52-emm28), 1314.3072 (ST39-emm4), and GCF_041430965.1 (emm-patternE-other). CDS features were extracted from Bakta-annotated GFF3 files, and locus tags were mapped to PanTA gene cluster identifiers using the gene presence/absence matrix with locus tag annotations. For each representative, a window of ±10 features around the anchor gene (sda1 or speC, respectively) was extracted, and display coordinates were computed relative to the anchor gene start codon (set to 0 kb). In cases where a prophage was integrated in the opposite chromosomal orientation relative to the other representatives (GCF_041430965.1 in Fig. 6), coordinates were reverse-complemented for direct visual comparison; the original genomic coordinates, display coordinates, and strand annotations are provided in **Table S9**. Genes were assigned to functional categories (phage structural, lysis, integration, DNA modification/replication, paratox, hyaluronidase, cargo, or chromosomal) based on Bakta product annotations.

#### Nested prophage analysis in genome 1314.4589

To characterize the tandem Sda1-encoding prophage duplication in ST28-emm1 strain 1314.4589 (**Fig. S6**), we identified the boundaries of the canonical prophage (Prophage 1: BENLNG_01456 to BENLNG_01545, ∼68 kb) and the nested prophage (Prophage 2: BENLNG_01470 to BENLNG_01504, ∼28 kb) from the Bakta-annotated GFF3 file. Protein sequences for all CDS within Prophage 1 (excluding the nested Prophage 2 region) and all CDS within Prophage 2 were extracted by translating the corresponding nucleotide sequences from the embedded genome FASTA. BLASTp (BLAST+ v2.17.0, default parameters, E-value threshold 1×10⁻⁵, max_target_seqs = 1) was used to identify the best Prophage 1 match for each Prophage 2 protein. The canonical sda1 (BENLNG_01457) and paralog sda1_00717 (BENLNG_01492) were additionally compared at the nucleotide level using BLASTn. Per-gene percent identity, alignment length, and functional annotations are reported in **Table S8**.

#### Isolation site classification and invasive isolation rate

Isolation sites were classified from the standardized source_cat metadata column, with the following additional rules. First, the seven ST63-emm77 strains originally tagged as “respiratory tract” in BV-BRC metadata, which derive from a Polish surveillance study of upper-respiratory pharyngitis isolates (51, 52), were reclassified as throat/pharyngeal. Second, strains with source_cat = “Clinical (Unspecified)” whose isolation_source field contained “normally sterile site” were classified as sterile-site isolates. Third, raw metadata fields (isolation_source, disease, isolation_comments) were scanned for explicit invasive-disease keywords (invasive, STSS, necrotizing fasciitis, bacteremia, sepsis, meningitis, cellulitis); strains matching these keywords but lacking a specific anatomical-site classification were assigned to “Invasive (unspecified).” Strains with source_cat = “Not Specified” and no invasive keywords, as well as non-anatomical categories (genital, urine, ear), were excluded from the denominator. Invasive isolation rate was computed as the percentage of strains from invasive sites (blood, sterile site, CSF, bone/joint, soft tissue) or with explicit invasive-disease labels among strains with available source metadata (**Table 1**).

## Data Availability

All analyses were performed on a dedicated server (128 CPU cores, 500 GB RAM) running Ubuntu 24. Key software versions: Bakta v1.12.0, PanTA (GitHub, April 2026), CD-HIT v4.8.1, BLAST+ v2.17.0, DIAMOND v2.1, emmtyper v0.2.2, CheckM v1.2.2, Mash v2.3, Python 3.12, scikit-learn v1.8.0, pandas v3, matplotlib v3.10.9, Plotly v6.6.0. The optNMF algorithm is implemented in the PyPhylon Python package (https://github.com/SBRG/pyphylon/). Genome assemblies are available from BV-BRC and NCBI under their original accession numbers. The Spyo-GENOMiCUS-400 genome compendium metadata, gene presence/absence matrices, binarized **L** and **A** matrices, and analysis notebooks are available at Zenodo (DOI: 10.5281/zenodo.20403113).

## Notes

https://doi.org/10.5281/zenodo.20403112

### Competing Interest Statement

The authors have declared no competing interest.

