## Supplementary Information for "Pangenomic decomposition reveals lineage-specific immune-evasion and virulence architectures in group A *Streptococcus*"

### Table Of Contents

**Supplementary Text** 3
Supplementary Note S1: Detailed composition of the S. pyogenes core-genome 3
S1.1 Functional composition of the omnipresent-core 3
S1.2 Composition and occupancy distribution of the soft-core 4
S1.3 Genes immediately below the strict-core threshold 4
S1.4 Two novel sub-lineage-specific structural variations in the core genome 5
S1.5 Evolutionary interpretation of the two novel core-genome findings 6
S1.6 Phenotypic predictions arising from the two novel core-genome findings 6
**Supplementary Tables** 7
**Supplementary Figures** 8
Supplementary Figure S1 8
Supplementary Figure S2 9
Supplementary Figure S3 10
Supplementary Figure S4 11
Supplementary Figure S5 12
Supplementary Figure S6 13
Supplementary Figure S7 14
**References** 16

### Supplementary Text

#### Supplementary Note S1: Detailed composition of the *S. pyogenes* core-genome

This Supplementary Note provides the per-functional-category and per-gene detail that supports the condensed core-genome characterization in the main Results. The full per-gene occupancy data are given in **Supplementary Table S1**; the functional-category breakdown is given in **Supplementary Table S2**.

##### S1.1 Functional composition of the omnipresent-core

The 747 genes of the *S. pyogenes* omnipresent-core partition across the standard COG functional categories as follows:

**Translation (140 genes; 18.7% of the omnipresent-core).** This is the largest single COG functional category (J), encompassing 44 of the 58 ribosomal proteins, all 20 aminoacyl-tRNA synthetase activities (some, like methionyl-tRNA ligase, distributed across two paralogous omnipresent clusters), the elongation factors EF-Tu (*tuf*) and EF-G (*fusA*), initiation factors IF-1, IF-2, and IF-3, and the rRNA modification machinery.

**Carbohydrate transport and metabolism plus energy production (99 genes; 13.3%; combining COG categories G and C).** The glycolytic pathway is anchored in the omnipresent-core via *pyk* (pyruvate kinase), *pfkA* (6-phosphofructokinase), and *gap* (glyceraldehyde-3-phosphate dehydrogenase). The remaining glycolytic enzymes – *pgk* (398/399), *eno* (395/399), *ldh* (398/399), and an *fbaA*-class aldolase – fall one genome below 100% occupancy and are therefore part of the soft-core rather than the omnipresent-core.

**DNA replication, recombination, and repair (35 omnipresent genes; COG L).** DNA replication is fully represented in the omnipresent-core: DnaB-helicase, DnaG primase, PolA, GyrA, ParC, and SSB are all present in 399/399 genomes.

**Nucleotide transport and metabolism (47 omnipresent genes; COG F).** This category contributes 47 omnipresent genes spanning the canonical purine and pyrimidine biosynthetic pathways.

**Posttranslational modification, protein turnover, chaperones (34 omnipresent genes; COG O).** The heat-shock and protein quality-control axis (GroES, DnaK, DnaJ, GrpE, ClpP, and ClpX) is fully represented in the omnipresent-core, consistent with its essentiality for viability under host-imposed thermal and oxidative stress.

**Regulatory genes.** The principal regulatory genes of the species are all omnipresent: CovR (response regulator controlling virulence-gene expression), CodY (GTP-sensing pleiotropic regulator), RelA ((p)ppGpp synthase/hydrolase mediating the stringent response), VicK (cell-wall metabolism histidine kinase), and LiaF (cell-envelope stress response).

**COG categories M (cell envelope; 40 omnipresent genes) and D (cell cycle / division; 12 omnipresent genes).** FtsZ, FtsA, DivIVA, the *mur* peptidoglycan biosynthesis operon, all six penicillin-binding-protein activities, and the canonical sortase A are present in every genome.

**Virulence-related genes.** Among classical *S. pyogenes* virulence factors, only the NADase *nga* is fully omnipresent. Every other classical *S. pyogenes* exotoxin or surface virulence factor occupies between 394 and 398 of 399 genomes.

**No COG assignment.** 41 of the 747 omnipresent-core genes (5.5%) lack a COG assignment, reflecting their lineage-specific origins outside the general bacterial proteome captured by COG. These include the canonical *S. pyogenes* NADase (*nga*) and several conserved hypothetical proteins.

#### S1.2 Composition and occupancy distribution of the soft-core

The 545 genes of the soft-core have a strongly right-skewed occupancy distribution: 302 of 545 (55%) are absent in only a single genome, and 496 of 545 (91%) are present in 99% or more of genomes. The soft-core therefore does not constitute a separate functional layer but rather completes pathways already established by the omnipresent-core, with a small fraction of genuinely lineage-variable genes at lower occupancy.

**Glycolysis completion.** Glycolysis is completed by *pgk* (398/399), *eno* (395/399), *ldh* (398/399), and an *fbaA*-class aldolase.

**Virulence factors.** The major *S. pyogenes* virulence factors are largely soft-core: *sagA-G* streptolysin S biosynthesis, *speB* cysteine protease, *ska* streptokinase, *scpA* C5a peptidase, *spd* streptodornase, and *cfa* CAMP factor each occupy between 394 and 397 of 399 genomes.

**Regulators.** The canonical CovS sensor kinase (397/399) and the RopB/Rgg quorum regulator (394/399) sit in the soft-core, complementing the omnipresent CovR response regulator.

**Cell-envelope decoration.** The *dlt* operon for D-alanyl-lipoteichoic-acid biosynthesis (394/399) sits in the soft-core.

**Transport.** A comparable pattern holds for the 54 soft-core transport genes: the omnipresent-core typically retains the ATP-binding subunit and one permease of a given ABC transporter, while the substrate-binding protein or the second permease falls into the soft-core.

#### S1.3 Genes immediately below the strict-core threshold

Several canonical *S. pyogenes* virulence factors and regulators occupy the immediate sub-core boundary at 82-97% occupancy. These are not threshold artefacts but biologically meaningful lineage-restriction patterns.

**Capsule biosynthesis operon (*hasA* 83%, *hasB* 86%, *hasC* 88%).** This pattern reflects the capsule-deficient phenotype of the M89 (ST101) lineage and other invasive *S. pyogenes* lineages, including *emm*4, *emm*22, *emm*28, and *emm*87.

**Streptolysin O (*slo*; 97%).** SLO is a near-universal cytolysin; the missing 3% reflects sequence divergence in a small subset of genomes rather than gene loss.

***mga* (97%).** The major virulence regulator *mga* shows a similar pattern, reflecting two divergent allelic forms (Mga-1 in the A-C *emm*-cluster and Mga-2 in the E/D *emm*-cluster) partitioned across major *S. pyogenes* lineages.

**Other lineage-restricted surface adhesins.** *scl1* (46%) and *fbaA* (12%) fall well below the core threshold and partition into the accessory-genome.

#### S1.4 Two novel sub-lineage-specific structural variations in the core genome

Beyond the established core-genome distributions described above, our analysis identified two sub-lineage-specific structural variations not previously emphasized in the *S. pyogenes* literature. Both are consistent with single ancestral events that have been clonally propagated through their respective sub-lineages [1,2] and predict loss-of-function phenotypes for affected isolates.

**Coordinated cassette substitution at the *cfa* locus in ST382-*emm*6.4.** All four ST382-*emm*6.4 genomes in the dataset carry a coordinated replacement at the *cfa* (CAMP factor) locus, in which a 5.6 kb reference segment present in the SF370 reference genome is replaced by a 3.05 kb cassette encoding an alternative gene complement. The replacement-block dimensions match exactly across all four genomes, with identical breakpoints and preserved flanking-gene synteny, consistent with a single ancestral homologous recombination event rather than four independent acquisitions. The cassette substitution eliminates the canonical *cfa* open reading frame while preserving the surrounding chromosomal architecture. Because CAMP factor contributes to co-hemolysis and host-cell membrane disruption in group A streptococci, this substitution predicts loss of CAMP-factor-mediated co-hemolytic activity in the ST382-*emm*6.4 sub-lineage (**Fig. S4; Table S3**). Further phenotypic implications are discussed in **Note S1.5**.

**Clonally propagated *puuD* pseudogene in ST15.** Five of the ten ST15 genomes in the dataset carry a *puuD* gene that has been split into two fragments (345 bp + 312 bp) separated by a 39-bp intergenic gap, producing a pseudogene incapable of encoding a full-length protein. The inter-fragment geometry (fragment sizes and gap length) is identical across all five affected genomes, consistent with a single ancestral nonsense or frameshift mutation rather than independent disruption events. The remaining five ST15 genomes retain an intact *puuD* open reading frame. PuuD (gamma-glutamyl-gamma-aminobutyrate hydrolase) catalyzes a step in the putrescine utilization pathway, and its disruption predicts loss of polyamine catabolism in the affected sub-lineage, potentially altering nitrogen metabolism and fitness under polyamine-rich conditions such as those encountered during tissue invasion (**Fig. S5; Table S4**). Further phenotypic implications are discussed in **Note S1.6**.

Both patterns are consistent with the broader observation that recombination and clonal propagation of structural mutations drive much of the within-lineage variation in *S. pyogenes* [1,2]. These findings illustrate how pangenome-level analysis at the core-accessory boundary can detect sub-lineage structural events that are invisible to single-gene or single-strain analyses.

#### S1.5 Evolutionary interpretation of the two novel core-genome findings

Both findings are consistent with single ancestral recombination or mutation events that have been clonally propagated through their respective sub-lineages. For ST382-emm6.4, replacement-block dimensions match exactly across all four genomes (5.6 kb reference segment replaced by a 3.05 kb cassette, with identical breakpoints and flanking-gene synteny preserved), consistent with a single ancestral homologous recombination event rather than independent acquisitions. For ST15, the inter-fragment geometry of the puuD split (345 bp + 39 bp gap + 312 bp) is identical across all five affected genomes, consistent with a single ancestral nonsense or frameshift mutation. Both patterns are consistent with the broader observation that recombination drives much of the within-lineage structural variation in S. pyogenes [1,2].

#### S1.6 Phenotypic predictions arising from the two novel core-genome findings

Both novel findings reported in the main Results carry concrete, testable phenotypic predictions for the affected sub-lineages:

**ST382-*emm*6.4 *cfa*-locus replacement.** ST382-*emm*6.4 isolates should:

- Lack CAMP-factor activity (loss of *cfa*).
- Lack polar-amino-acid uptake of histidine, glutamine, and arginine (loss of *hisJ*, *glnQ*, *hisM*).
- Lack the function of the *fieF* cation-efflux pump and the *phnA* protein.
- Gain cobalt/zinc/cadmium tolerance (acquisition of the cobalt-zinc-cadmium efflux pump).
- Gain mannitol-utilization capacity (acquisition of the mannitol-specific PTS transporter).

**ST15 *puuD* pseudogenisation.** The five ST15 *puuD*-pseudogene isolates should lack γ-glutamyl-γ-aminobutyrate hydrolase activity in their polyamine-catabolism pathway, with the upstream and downstream metabolic intermediates accumulating accordingly.

These predictions are testable in extant ST382 and ST15 isolates from public strain collections.

### Supplementary Tables

**Supplementary Tables.** Owing to size, the Supplementary Tables for this manuscript are deposited on Zenodo at **DOI: 10.5281/zenodo.20403112**.

- **Table S1** — Per-gene occupancy and COG functional-category assignment for all 1,292 core genes of Spyo-GENOMiCUS-400 (1,292 rows × 17 columns).
- **Table S2** — COG functional-category breakdown of the omnipresent-core and soft-core layers (24 rows × 11 columns).
- **Table S3** — Per-genome locus_tag identifiers, coordinates, strands, lengths, and product annotations for the *cfa*-locus region in the five genomes of Supplementary Figure S4 (105 rows × 13 columns).
- **Table S4** — Per-genome locus_tag identifiers, coordinates, strands, lengths, and product annotations for the *puuD*-locus region in the seven genomes of Supplementary Figure S5 (147 rows × 13 columns).
- **Table S5** — Per-phylon binary concordance with established classification systems
- **Table S6** — Global multi-class concordance against established classification systems
- **Table S7** — Enrichment analysis of the 27-gene Sda1 prophage core across Sda1+ vs Sda1− *S. pyogenes* phylons. For each gene cluster, the table provides the PanTA gene cluster ID, Bakta product annotation, assigned functional module, total carriers across all 399 genomes, carriage in Sda1+ phylons (n=121 strains), carriage in non-Sda1+ phylons (n=278 strains), log-odds ratio (Haldane-Anscombe corrected), Fisher's exact test p-value (one-tailed, alternative=greater), Benjamini-Hochberg-corrected q-value, and binary enrichment call at FDR < 0.05 (27 rows × 12 columns).
- **Table S8** — BLASTp comparison of all 35 Prophage 2 (nested) proteins against their Prophage 1 (canonical) counterparts in ST28-emm1 strain 1314.4589. For each paralog pair, the table provides the Prophage 2 locus tag, Prophage 1 locus tag, percent protein identity, alignment length, number of mismatches, query length, subject length, query coverage, E-value, and Prophage 2 product annotation (35 rows × 10 columns).
- **Table S9** — Per-genome locus_tag identifiers, coordinates, strands, lengths, and product annotations for the *speC/mf2* synteny windows in the three representative genomes of **Fig. 6** (63 rows × 13 columns).

### Supplementary Figures

#### Supplementary Figure S1

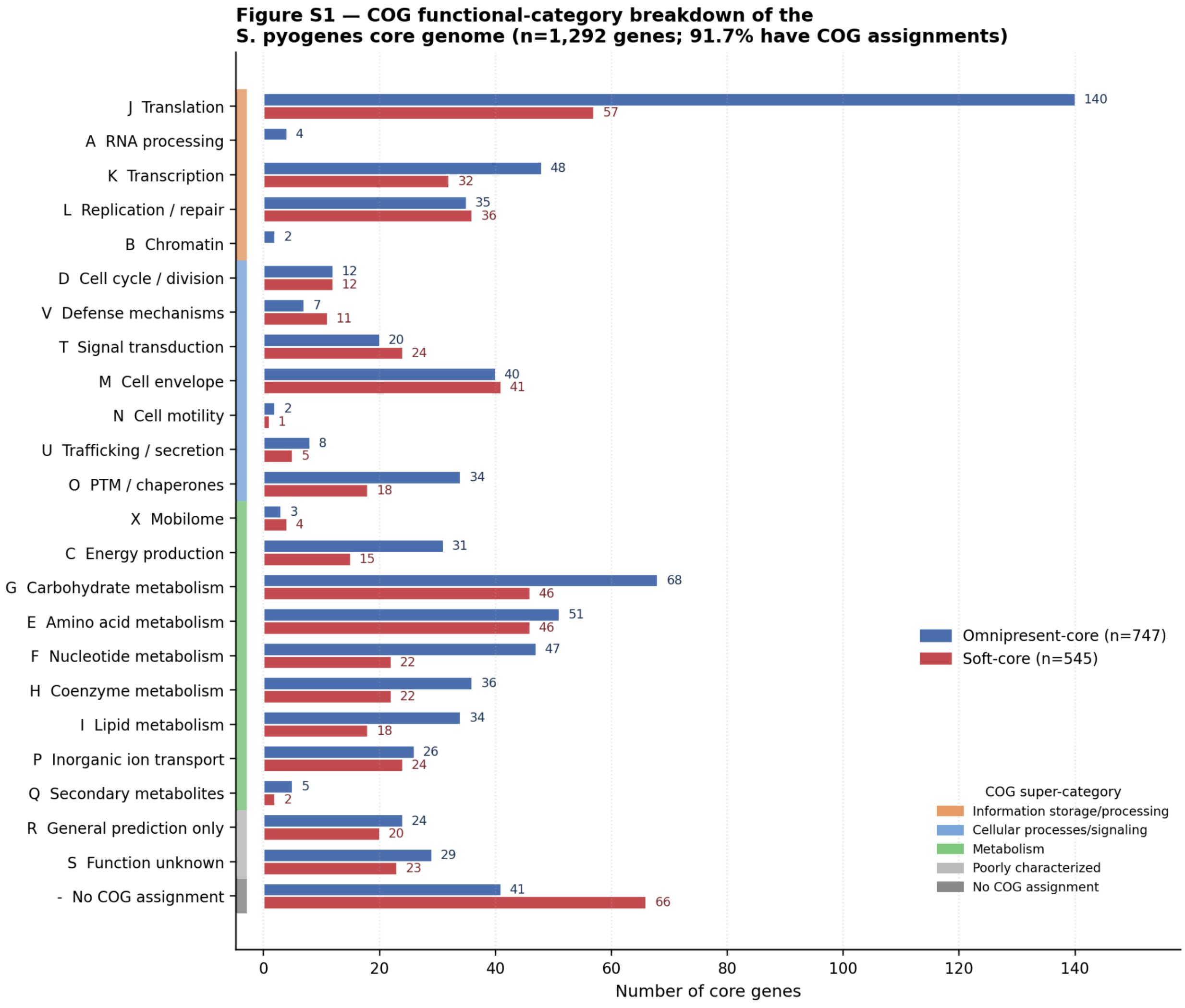


***Supplementary Figure S1: COG functional-category breakdown of the S. pyogenes core genome (n = 1,292 genes), partitioned into omnipresent-core (n = 747) and soft-core (n = 545).*** *Categories follow the canonical COG ordering grouped into the four COG super-categories (information storage and processing, cellular processes and signaling, metabolism, poorly characterized). 41 omnipresent-core and 66 soft-core genes lack a COG assignment, reflecting their lineage-specific origins outside the general bacterial proteome captured by COG.*

### Supplementary Figure S2

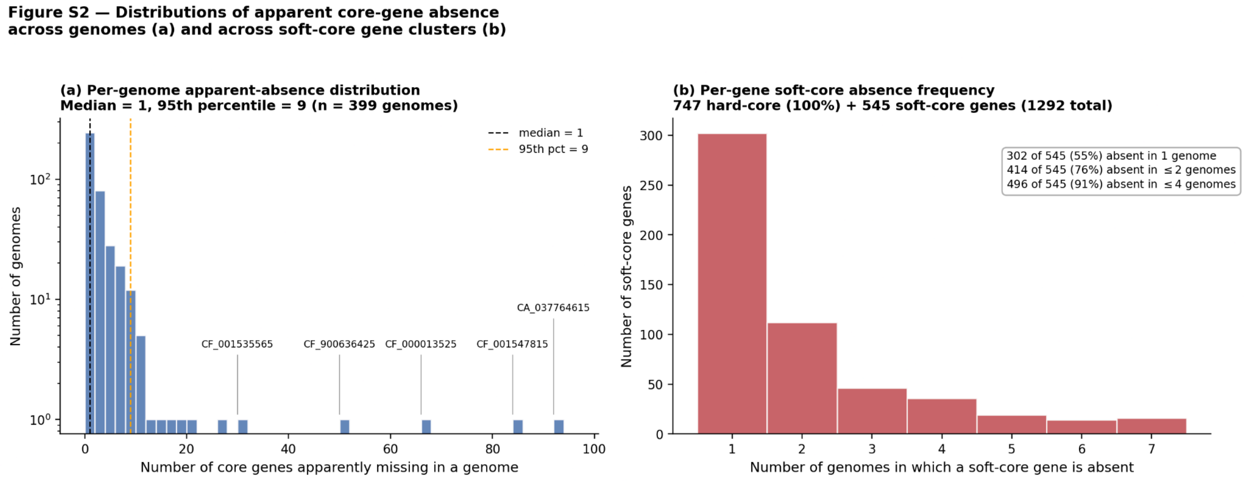


***Supplementary Figure S2: Distributions of apparent core-gene absence across genomes (a) and across soft-core gene clusters (b). Panel (a): the median S. pyogenes genome carries 1 missing core gene;*** *the right tail of 5 genomes carrying 30-92 apparent absences reflects sequence divergence routing canonical genes into separate PanTA gene clusters rather than true gene loss.* ***Panel (b): of the 545 soft-core genes, 91% are absent in 4 or fewer genomes, supporting the conclusion that the soft-core completes pathways established by the omnipresent-core rather than constituting a separate functional layer.***

### Supplementary Figure S3

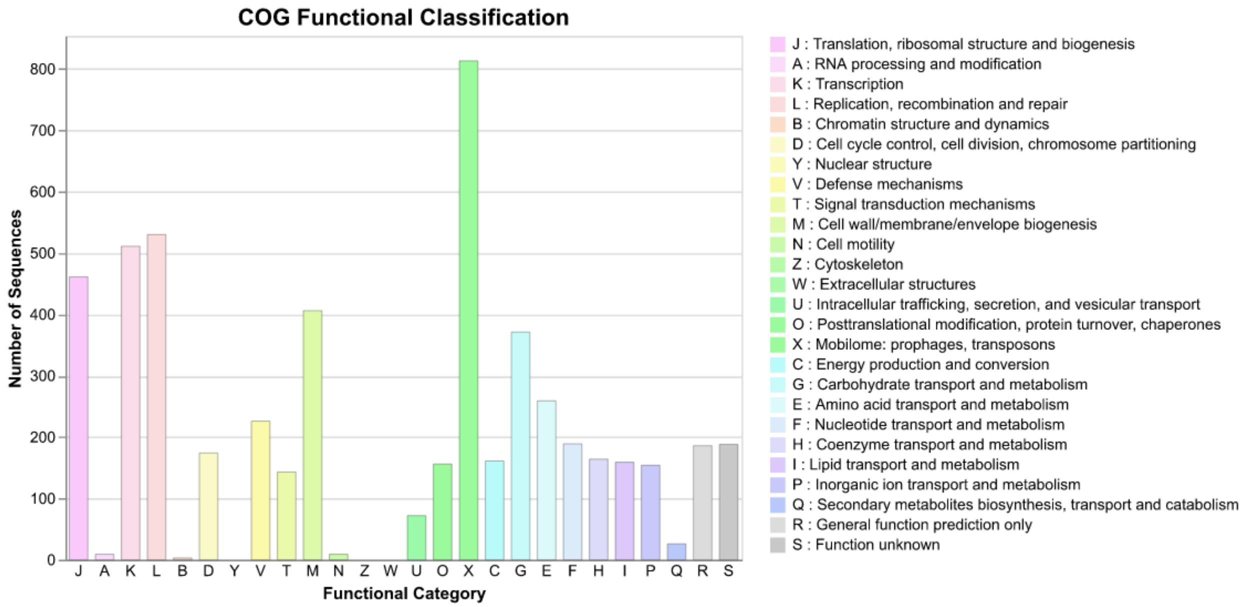


***Supplementary Figure S3: COG functional-category breakdown of the full S. pyogenes pangenome (5,370 COG-assigned genes from the 9,700-cluster pangenome), generated by COGclassifier with default settings.*** *Categories follow the canonical COG ordering. The dominant Mobilome category (X; 813 genes) reflects the prophage and transposon content of the S. pyogenes accessory genome.*

### Supplementary Figure S4

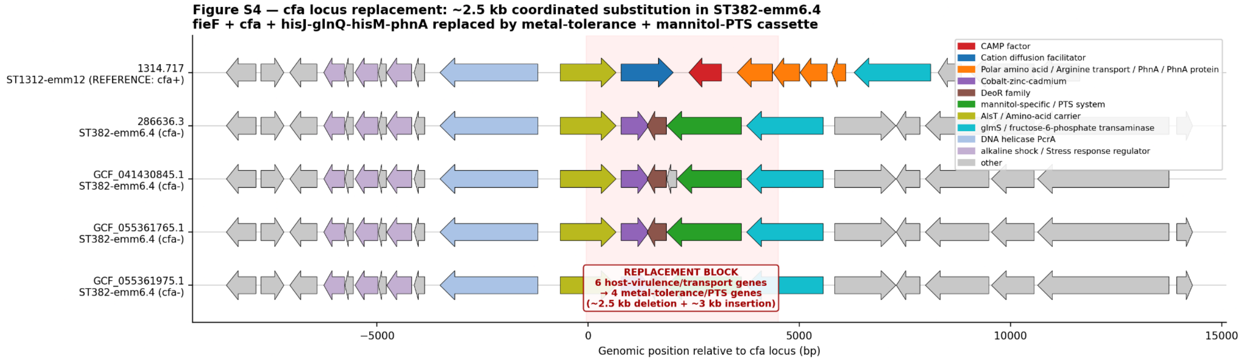


***Figure S4. The ST382-emm6.4 cfa-locus replacement.*** *Synteny of the cfa-locus region in the four ST382-emm6.4 genomes of Spyo-GENOMiCUS-400 (rows 2-5) compared to the reference genome 1314.717 (ST1312-*emm12*, cfa+; row 1). Each row shows ±10 features around the AlsT anchor on the genome's primary contig, with arrows indicating gene direction and color indicating functional class. The reference genome carries six contiguous host-virulence and polar-amino-acid-transport genes between AlsT (yellow) and* glmS *(cyan): a cation-efflux pump (*fieF*; blue), the CAMP-factor pore-forming toxin (*cfa*; red), and a four-gene polar-amino-acid uptake module (*hisJ, glnQ, hisM, phnA*; orange). In all four ST382-emm6.4 genomes, these six genes have been replaced (red-shaded region) by a different cassette encoding a cobalt-zinc-cadmium efflux pump (purple), a DeoR-family transcriptional regulator (brown), and a mannitol-specific phosphotransferase-system (PTS) transporter (green). Replacement-block dimensions match across all four ST382-*emm6.4 *genomes, with the replaced segment 2,546 bp shorter than the segment it replaces. Flanking-gene synteny (grey and pastel-colored arrows beyond the highlighted region) is preserved in all five genomes, indicating that the substitution is locally restricted to the* cfa *locus. Per-genome locus_tag identifiers, coordinates, and product annotations are provided in Supplementary Table S3.*

### Supplementary Figure S5

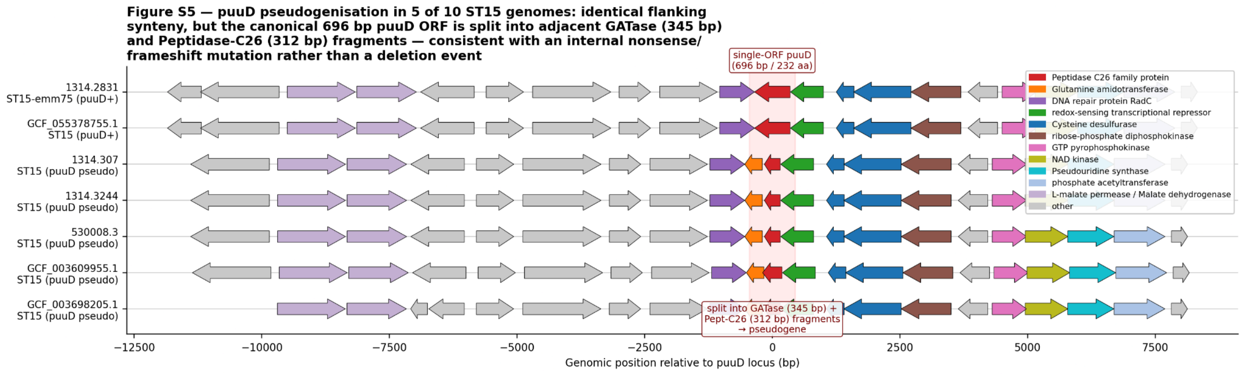


***Figure S5. The ST15 puuD pseudogenisation.*** *Synteny of the* puuD*-locus region in the seven ST15 genomes inspected here: two* puuD+ *reference genomes (1314.2831 and GCF_055378755.1; rows 1-2) and five* puuD*-pseudogene genomes (rows 3-7). Each row shows ±10 features around the Peptidase C26 anchor on the genome's primary contig, with arrows indicating gene direction and color indicating functional class. The reference genomes carry an intact 696 bp* puuD *coding sequence (red, central red-shaded region) flanked upstream by* radC *(purple, "DNA repair protein RadC") and downstream by rex (green, "redox-sensing transcriptional repressor"), cysteine desulfurase (blue), and the* yjbM */ NAD-kinase / pseudouridine-synthase /* pta *gene cluster. In the five* puuD*-pseudogene genomes, the canonical* puuD *coding sequence is replaced by two adjacent shorter open reading frames: a 345 bp upstream fragment retaining the N-terminal GATase-1 catalytic domain (orange, "Glutamine amidotransferase") and a 312 bp downstream fragment retaining the C-terminal Peptidase C26 domain (red), separated by a 39 bp intergenic gap. The combined extent of the two fragments matches the length of the intact* puuD *coding sequence, and the chromosomal neighborhood (flanking genes upstream and downstream) is preserved across all seven genomes. The locus in GCF_003698205.1 is shown flipped relative to its native chromosomal orientation to align visually with the other genomes; the underlying gene order is identical. Per-genome locus_tag identifiers, coordinates, and product annotations are provided in Supplementary Table S4.*

### Supplementary Figure S6

**A nested Sda1-encoding prophage in strain 1314.4539 indicates recent intra-genomic duplication of the canonical Sda1-encoding prophage**

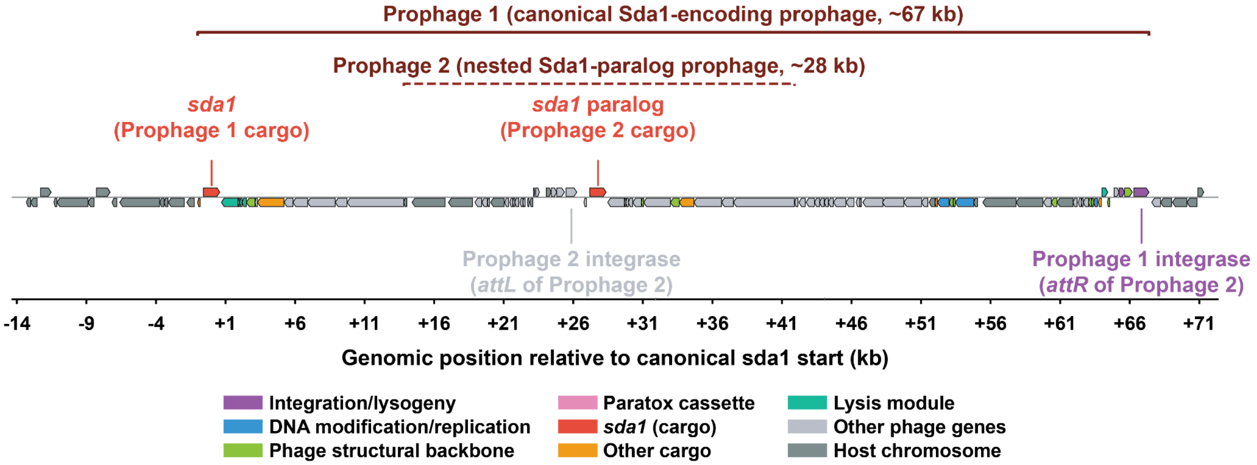


***Supplementary Figure S6: A nested Sda1-encoding prophage in ST28-emm1 strain 1314.4589.*** *Synteny of the double-prophage region in strain 1314.4589, which carries two tandemly integrated Sda1-encoding prophages on the same contig. Prophage 1 (canonical, ~68 kb; solid bracket) spans from the paratox gene (BENLNG_01456) to the integrase (BENLNG_01545) and contains 25 of the 27 Sda1+ shared-exclusive genes (color-coded by functional module as in Fig. 5). Prophage 2 (nested, ~28 kb; dashed bracket) is inserted within Prophage 1 and carries its own integrase (BENLNG_01490), Cro/CI lysogeny regulator, and a complete copy of the sda1 → amidase → holin → hyaluronidase → tail cargo cassette. The sda1 paralog (BENLNG_01492, PanTA cluster sda1_00717; dark red) is 100% identical to the canonical sda1 (BENLNG_01457) at both nucleotide (1,173/1,173 bp) and protein (390/390 aa) level. BLASTp comparison of all 35 Prophage 2 genes against Prophage 1 shows 27 of 35 pairs at 100% protein identity (mean 99.4%, range 87.8-100%;* ***Table S8****), with the greatest divergence in the Cro/CI regulator (87.8%) and integrase (98.7%) — the genes expected to diverge first for independent lysogenic maintenance of the nested element. Coordinates are shown relative to the canonical sda1 start codon. Per-gene locus_tag identifiers, percent identity, and alignment statistics are provided in Table S8.*

### Supplementary Figure S7

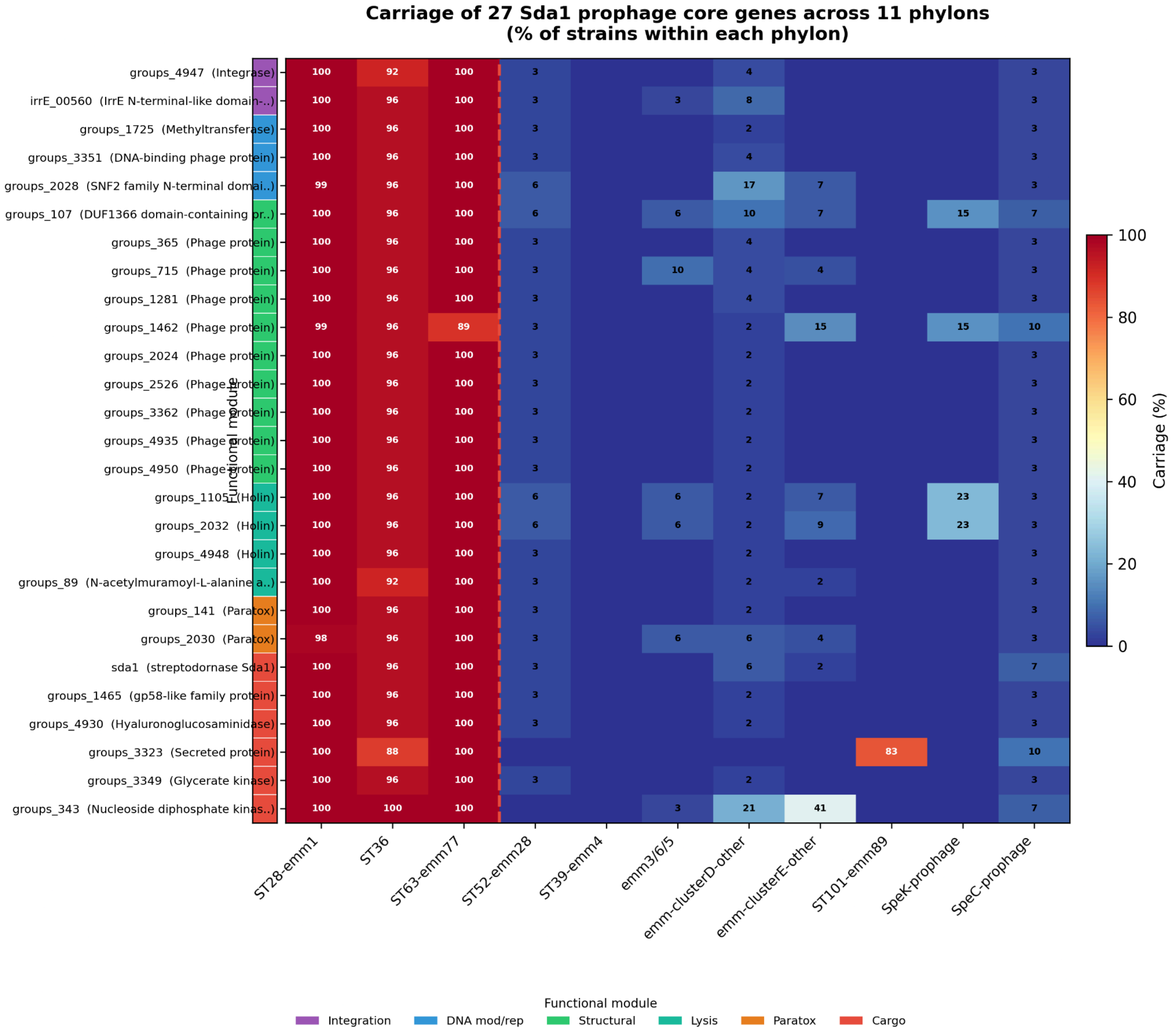


***Supplementary Figure S7: Carriage of the 27-gene Sda1 prophage core across all 11 S. pyogenes phylons.*** *Each cell shows the percentage of strains within a phylon that carry the indicated gene cluster (from the PanTA gene presence/absence matrix, n = 399 genomes). Genes (rows) are ordered by functional module within the canonical temperate-prophage architecture: integration (purple), DNA modification/replication (blue), phage structural backbone (green), lysis (teal), paratox cassette (orange), and accessory virulence cargo (red). Phylons (columns) are ordered with the three Sda1+ phylons (ST28-emm1, ST36, ST63-emm77) at left, separated from the six non-Sda1 taxonomic phylons and two mobilons by a dashed red line. The 25 in-prophage genes show tight co-segregation with the Sda1+ phylons (median carriage 99.2%) and near-zero background in non-Sda1 phylons (median 4.0%). Two genes at distant chromosomal positions — groups_343 (nucleoside diphosphate kinase, 22% in emm-clusterD-other) and groups_3323 (secreted protein, 83% in emm-clusterE-other) — show*

*elevated background outside the Sda1+ phylons, consistent with independent homologs or functions outside the prophage context. Per-gene enrichment statistics (Fisher's exact test, BH-corrected q-values) are provided in* ***Table S7****.*
